# BlpC-mediated selfish program leads to rapid loss of *Streptococcus pneumoniae* clonal diversity during infection

**DOI:** 10.1101/2022.08.05.500807

**Authors:** Surya D. Aggarwal, John A. Lees, Nathan T. Jacobs, Gavyn Chern Wei Bee, Annie R. Abruzzo, Jeffrey N. Weiser

## Abstract

Chromosomal barcoding and high-throughput sequencing were used to investigate the population dynamics of *Streptococcus pneumoniae*. During infant mouse colonization, >35-fold reduction in diversity and expansion of a single clonal lineage was observed within 1 day post-inoculation. This loss of diversity was not due to immune factors, host microbiota or exclusively because of genetic drift. Rather, it required the expression of *blp* bacteriocins induced by the BlpC-quorum sensing pheromone. This points towards the role of intra-strain competition whereby the subpopulation reaching a quorum eliminates others that have yet to activate the *blp* locus. We show that this loss of diversity also restricts the number of unique clones that could establish colonization during transmission between hosts. Moreover, we show that genetic variation in the *blp* locus is associated with transmissibility in the human population. We posit this is due to its importance in clonal selection and its role as a selfish element.

## INTRODUCTION

An important step in the lifecycle of opportunistic pathogens includes the successful colonization of a host. During this stage, microbes adapt to the host environment by altering their gene expression and subverting host defenses. Competitive success during host colonization, therefore, shapes the ecology of infecting microbes, and influences their pathogenesis and ability to transmit between hosts – influencing their population-wide success (Weiser et al., 2018).

While numerous studies have characterized specific factors important for colonization and transmission, little is known about the population dynamics of microbes *in vivo* (D’Mello et al., 2020; Matthews et al., 2021; van Opijnen and Camilli, 2012; Rowe et al., 2019; Zafar et al., 2019). Routinely used readouts such as population size do not provide the molecular resolution required for mechanistic insight into bacterial interactions during initial stages of infection. Technological advances have enabled such investigations through affordable deep-sequencing of chromosomally-barcoded bacteria and allowed for exploration of population bottleneck sizes during disease (Bachta et al., 2020; Fiebig et al., 2021; Hullahalli and Waldor, 2021; Hullahalli et al., 2021; Jasinska et al., 2020; Martin et al., 2017; Vasquez et al., 2021). However, these methods have not yet been used to study the entire infection lifecycle, and hence there is still little knowledge regarding the processes that shape population structure during colonization and transmission.

Using chromosomally-barcoded bacteria, we investigate the population dynamics of *Streptococcus pneumoniae* (*Spn*) during colonization and transmission. *Spn* is a common nasopharyngeal colonizer and a leading cause of disease following transit to normally sterile host sites. To study its pathogenic lifecycle, an animal model that recapitulates key features of human *Spn* pathogenesis is critical. Infant mice fulfill this need as they, like young children, exhibit high susceptibility to infection, prolonged colonization duration and enhanced propensity for transmission (Kono et al., 2016; Zafar et al., 2016). Using this animal model to investigate *Spn* population dynamics during colonization, we demonstrated a rapid loss in diversity of isogenic *Spn* clones over this period, which consequently impacted population structure during transmission. This loss of diversity was attributable to quorum sensing-dependent expression of bacteriocins, which we propose act selfishly to promote their own success in the *Spn* population.

## RESULTS

### Construction of a diverse DNA-barcoded *Spn* library

To evaluate the population structure of *Spn* and study the trajectory of multiple clonal lineages during infection, we constructed a DNA-barcoded library of a *Spn* 23F strain (**Figs 1A, S1**). A plasmid library containing short nucleotide sequences (barcodes), which tagged each clone with a distinct molecular identity, was transformed into the 23F strain (**Fig S1**). The resulting *Spn* 23F library consisted of 2,764 uniquely barcoded clones that are isogenic, differentiated only by the nucleotide sequence of their respective barcode (**Fig S1**). The barcode distribution of this *Spn* library followed a slightly over-dispersed Poisson distribution, with the most abundant clone in the library present at a frequency of 0.65%, suggesting a well-mixed distribution of uniquely barcoded clones. The abundance distribution of these clones remained unchanged upon *in vitro* passaging confirming stability of the barcoded *Spn* library (**Fig S2**).

**Fig 1:**
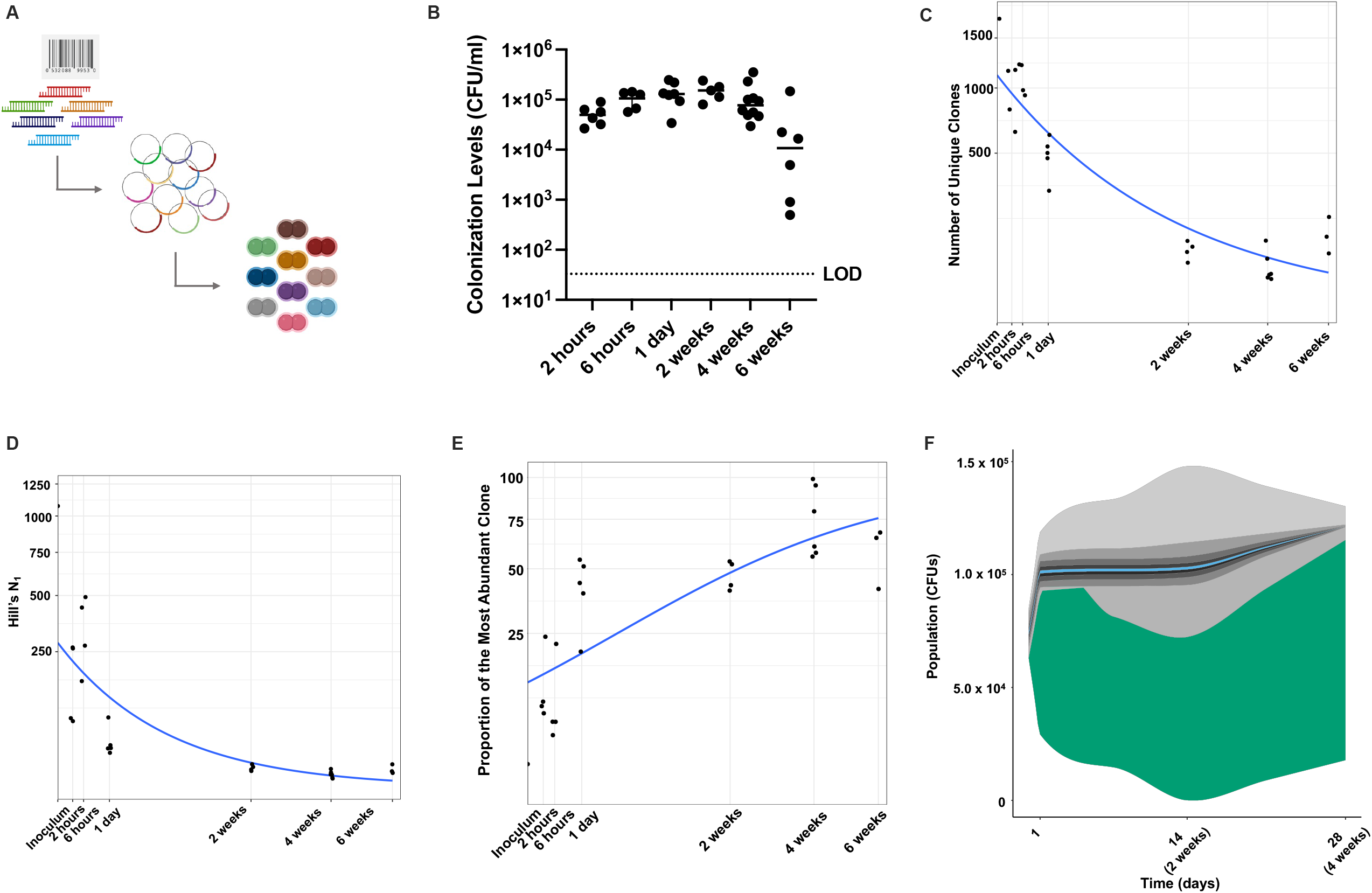
Rapid loss of within-host clonal diversity of *Spn* during nasopharyngeal colonization. (A) Schematic denoting the workflow of constructing *Spn* molecularly-barcoded clonal library. (B) Colonization levels of *Spn* library in retro-tracheal lavages from infant pups over time. Pups were infected at days 3-4 of age. LOD: Limit of Detection. (C-E) Diversity of the *Spn* library decreases over time as measured by (C) number of unique clones present (*P*<10^−5^ by quasipoisson regression), (D) Hill’s N_1_ diversity coefficient (*P*<10^−5^ by Poisson regression), and (E) proportion of the most abundant clone (*P*=1⍰10^−4^ by quasibinomial regression). *P*-values were calculated using a generalized linear model, and fitted trend is shown in blue line. (B-E) Each data point represents an individual animal. (F) Illustrative dynamics of top ten barcodes over time. Total height represents measured CFU/ml, and each band shows the proportion of the community made up by the clones, ranked by final frequency. Clone highlighted in blue is the most common barcode in the inoculum while the one highlighted in green is most abundant at 4 weeks.

### Rapid loss of *Spn* clonal diversity during colonization

Next, we investigated how the population structure of *Spn* changes during colonization in an infant mouse model. Upon inoculation with the *Spn* 23F library, the bacterial colonization density increased steadily, until reaching carrying capacity 1 day post-inoculation (**Fig 1B**). Colonization levels remained unchanged for up to 4 weeks. Over 50% of the clones present in the inoculum were successful in establishing colonization as early as 2-and 6 hours post-inoculation (**Fig 1C**). However, there was a sharp decline in the number of unique clones present during the course of colonization (**Fig 1C**). At 1 day post-inoculation, only 28.2% of the total clones were recovered from the mice, while the corresponding number at 2 weeks was 4.3%. The clonal diversity, measured by Hill’s N_1_ coefficient, also decreased rapidly to 2.77% of the inoculum at 1 day post-inoculation (**Fig 1D**). This diversity continued to decrease over time, with Hill’s N_1_ decreasing to 0.49% and 0.23% at 2 and 4 weeks, respectively, relative to the inoculum. The mean richness of the most abundant clone in the population increased from 0.65% in the inoculum to 41.6% at 1 day and 74.5% at 4 weeks post-inoculation (**Fig 1E, F**).

### Neither clonal abundance, genetic drift, nor *in vivo* adaptation explains the observed loss in diversity

We investigated whether success of a given clone during colonization is a function of the abundance of its founder in the inoculum. While the clone that was most abundant in the inoculum was detected in pups at 1 day post-inoculation, it was rare (<5% abundance) in each individual pup, and not detected in any pup at 2 or 4 weeks post-inoculation (**Fig 2A**). These data suggested that the clonal success of certain lineages was not necessarily a factor of their abundance in the inoculum. The identities of the most abundant clones detected in individual pups at each time points were distinct, as evidence of the barcodes being neutral (**Fig S3**).

**Fig 2:**
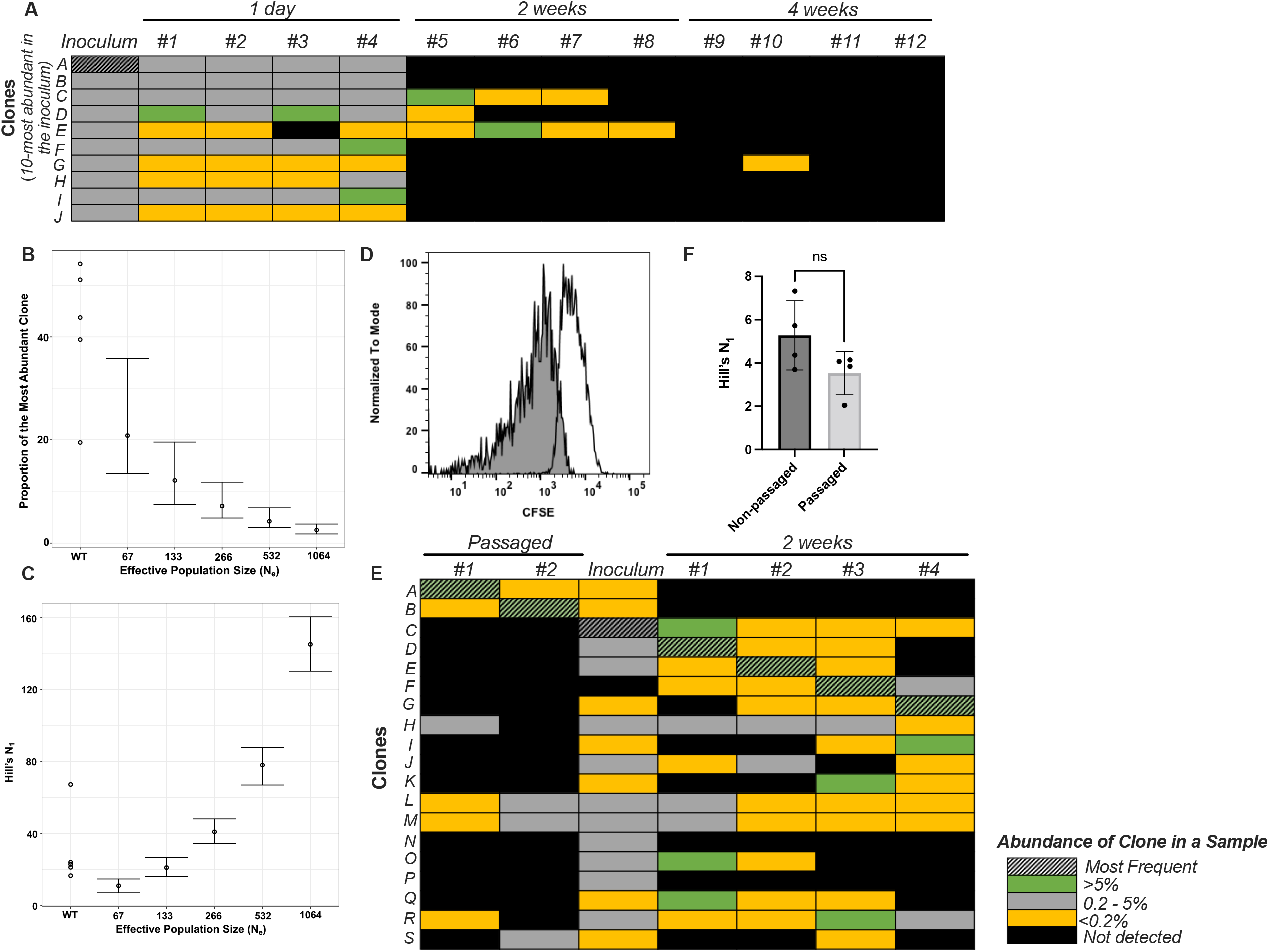
Random fluctuations, genetic drift and *in vivo* adaptation are not responsible for the observed loss in diversity. (A) Abundance of the clone in the inoculum is not an indicator of its success during colonization. Heat map denotes the abundance of individual clones belonging to the *Spn* library either in inoculum and individual infant pups (#1-12) at different time points. The clones selected for visualization were the 10-most abundant clones in the inoculum. The most abundant clone in the inoculum (clone A, hatched) is not detected in the populations found colonizing URT of pups at 2 and 4 weeks post inoculation. (B, C) Observed loss in diversity during early colonization (1 day post-inoculation) is unlikely to be attributed solely to genetic drift. WT denotes the observed values while the remaining points denote the expected values at different effective population sizes if loss in diversity was solely occurring due to genetic drift. Points show the mean from 300 repeats, and bars the 95% quantile. (D) Flow plot demonstrating a majority of the *Spn* population replicates *in vivo*. Blank curve denotes inoculum and the shaded curve represents population recovered at 19 hours post-inoculation. (E, F) *In vivo* adaptation is not responsible for the observed loss in diversity at 2 weeks post-inoculation. (E) Heat map denotes the abundance of individual clones belonging to the *Spn* library in the *in vivo* passaged populations, mixed inoculum and populations recovered from infant pups. Clones A and B that were most abundant in the passaged populations (hatched) were not detected from the URT of mice after 2 weeks following the re-inoculation of passaged populations into the pups. (F) Graph represents Hill’s N_1_ diversity coefficient when infected with non-passaged and *in vivo* passaged libraries. ‘ns’ denotes not statistically-significant by Mann-Whitney test.

We then investigated the contribution of genetic drift to the observed loss in diversity during colonization. Using Wright-Fisher simulations, we estimated the baseline expectation of clonal abundance that was influenced only by genetic drift at 1 day post-inoculation. The contribution of genetic drift depends on the dynamics of the actively replicating population, or effective population size, N_e_. Previous work has calculated a fairly low point estimate of 133 for N_e_ of *Spn* during nasal carriage in adult mice (Li et al., 2013). By modeling the experimental data obtained at 6 hours post-inoculation and at this low N_e_, the expected frequency of the most abundant clone and the Hill’s N_1_ values are significantly different than the observed values (**Figs 2B, C**). Given their higher carrying capacity of colonizing *Spn*, we postulated that infants may allow for larger N_e_ values relative to adult mice (Kono et al., 2016; Siegel et al., 2015). We used a previously described carboxyfluorescein diacetate succinimidyl ester (CFSE) dilution assay to determine what proportion of *Spn* are undergoing replication *in vivo* (Abruzzo et al., 2022; Siegel et al., 2014). We observed a pronounced reduction in the fluorescence intensity of CFSE among colonizing *Spn* at 19 hours post-inoculation, suggesting that a vast majority of cells have undergone division (**Figs 2D, S4**). 95.3% of the cells (10^5^ CFU/ml) present in the colonizing population had undergone at least one division by this time point. These data indicate that the N_e_ of *Spn* colonizing infant mice during these early time points may be significantly larger than 133. Combined, these data indicated that genetic drift alone was not the sole cause of lost diversity during colonization.

To test the possibility that *in vivo* adaptation of clones is responsible for their success, we challenged pups with a previously animal*-*passaged *Spn* library that were mostly dominated by single clones (∼95% and ∼99% abundance at 4 weeks, **Fig 1E**). Before inoculation, these animal-passaged populations were mixed with a non-passaged *Spn* library so that the overall abundance of the most abundant clones from the passaged populations was equivalent to the median abundance of clones in the non-passaged library. Of note, two weeks after this mixed library inoculation, the most abundant clones in these pups originated from the non-passaged library, not the previously *in vivo*-passaged populations (**Fig 2E**). Further, the diversity of the non-passaged and passaged libraries at 2 weeks was not statistically different (**Fig 2F**). These data excluded the possibility that mutations arose during *in vivo* passaging that conferred a fitness advantage to a particular clone.

### Host factors and co-infection do not contribute to loss of diversity during colonization

To explain the observed changes in population structure, our population genetics calculations allowed for two possibilities – either the most abundant clone replicated at a rate about twice as fast as other clones in the population, or was cleared at a rate approximately two-times slower (**Fig S5**).

Innate (TLR2 and IL-1 signaling) and adaptive (IL-17 signaling) immune factors have previously been shown to influence persistence by aiding in clearance of colonizing *Spn* (Kuipers et al., 2018; Lu et al., 2008; van Rossum et al., 2005; Zhang et al., 2009). We tested whether these factors also influenced the population structure of *Spn* at 4 weeks post-inoculation. There was no change observed in the diversity of colonizing *Spn* in either *Tlr2*^*-/-*^, *Il1r*^*-/-*^ or *Il17ra*^*-/-*^ mice (**Figs 3A–C**). The increased persistence of *Spn* was evident in *Il17ra*^*-/-*^ mice even at 4 weeks post-inoculation (**Fig 3D**).

**Fig 3:**
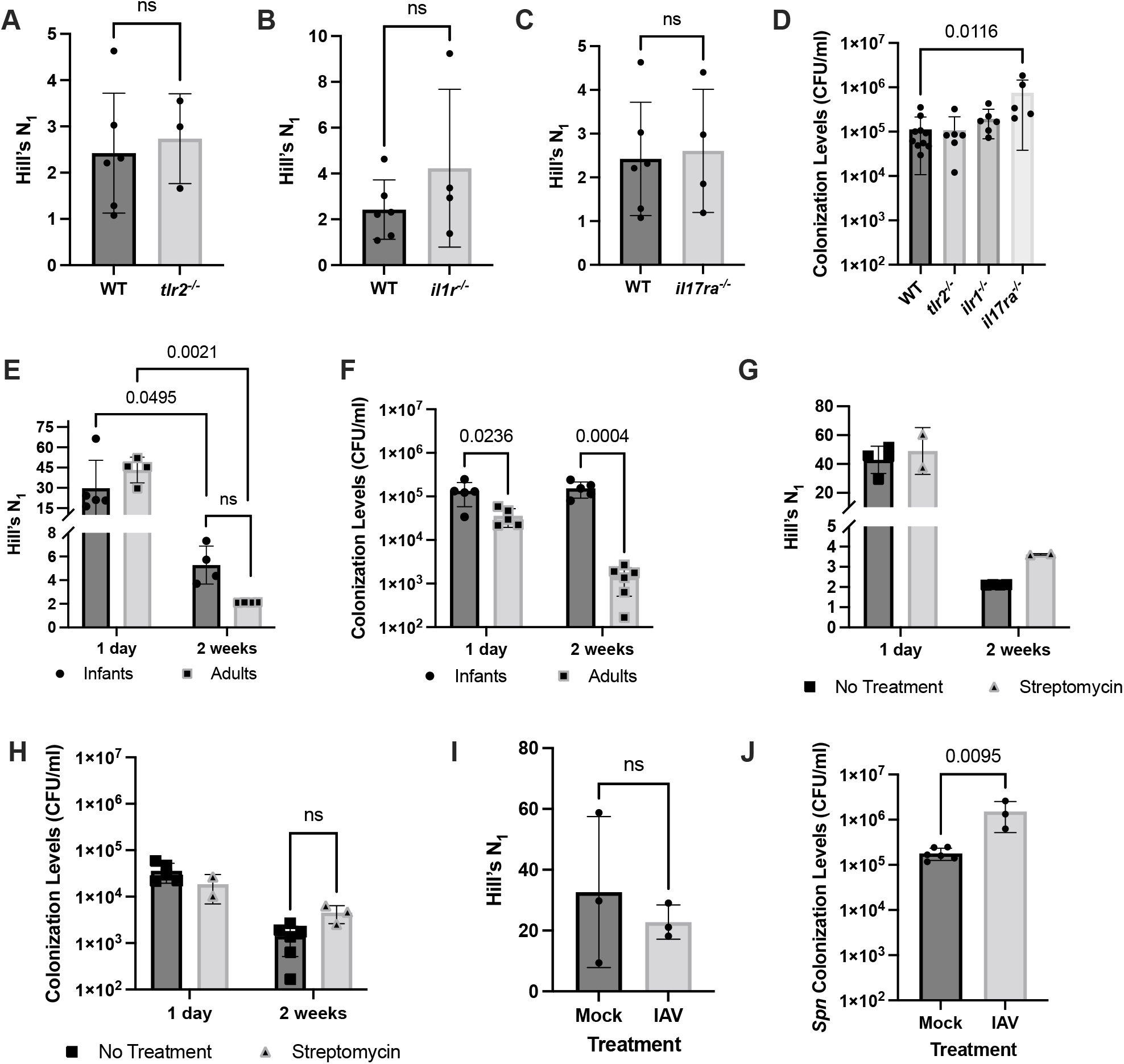
Host factors do not contribute to the observed loss of diversity. (A-D) Signaling by innate and adaptive immune factors (A) TLR2, (B) IL1R, and (C) IL17Ra that contribute to *Spn* clearance do not aid in loss of diversity at 4 weeks post inoculation. (D) Corresponding *Spn* colonization levels in retro-tracheal lavages from WT, *Tlr2*^*-/-*^, *Il1r*^*-/-*^, and *Il17ra*^*-/-*^ infant pups. *P*-values were calculated by Kruskal-Wallis test followed by Dunn’s correction for multiple comparisons. (E, F) Age has no effect on influencing *Spn* clonal diversity. Comparison of (E) Hill’s N_1_ diversity coefficient, and (F) colonization levels in infants and adults at 1 day and 2 weeks post-inoculation. *P*-values were calculated by 2-way ANOVA followed by Tukey’s multiple comparison test. (G, H) Depletion of host microbiota by streptomycin treatment did not influence *Spn* (G) Hill’s N_1_ diversity coefficient, or (H) colonization levels. (I, J) Influenza A virus (IAV) co-infection does not alter *Spn* population structure. Comparison of (I) Hill’s N_1_ diversity coefficient, and (J) *Spn* colonization levels in IAV co-infected pups relative to mock treatment group 10 days following *Spn* infection. *P*-values were calculated by Student’s *t-*test.

Age of the mice also did not influence the population structure of the colonizing *Spn* population. Similar to that in infant mice, we also observed a rapid loss of clonal diversity in adult mice (**Fig 3E**). As expected, *Spn* colonized infant mice at a higher density relative to adult mice (**Fig 3F**). These data also showed that increased colonization density did not correlate with higher diversity.

We investigated the hypothesis that prior occupation of the host niche by its resident microbiota restricts colonization by *Spn* and in turn, limits clonal diversity. The resident microbial community was disrupted by treating adult mice with streptomycin, previously shown to deplete the upper respiratory tract microbiota (Shen et al., 2019). Depletion of resident microbiota by streptomycin treatment did not significantly alter the colonization density or the clonal diversity of *Spn* (**Figs 3G, H**). While co-infection with influenza A virus (IAV) increased *Spn* colonization levels, such co-infection similarly had a negligible effect on diversity (**Fig 3I, J**).

### BlpC-mediated signaling is responsible for loss in *Spn* diversity during colonization

We then investigated if the observed decrease in diversity could be attributed to a bacterial factor. Given that all colonizing *Spn* were isogenic, differing only by unique barcodes, we postulated that quorum sensing-induced transcriptional variations in the colonizing population may affect population structure. Previous work has demonstrated that competence-induced activation of fratricidal molecules allows earlier colonizing bacteria to competitively exclude later arriving isogenic bacteria (Shen et al., 2019). We tested whether these fratricidal molecules (CibAB and CbpD) were also responsible for altering the *Spn* population structure during colonization by constructing a diverse Δ*cibAB*Δ*cbpD* barcoded-library (**Fig S6A**). No difference was observed in either the colonization density or clonal diversity of WT versus Δ*cibAB*Δ*cbpD* strains 1 day post-inoculation (**Figs 4A, B**). Thus, these fratricidal molecules that are important for *in vivo* competition during early hours following colonization (Abruzzo et al., 2022; Shen et al., 2019), do not lead to the observed decrease in diversity of *Spn*.

**Fig 4:**
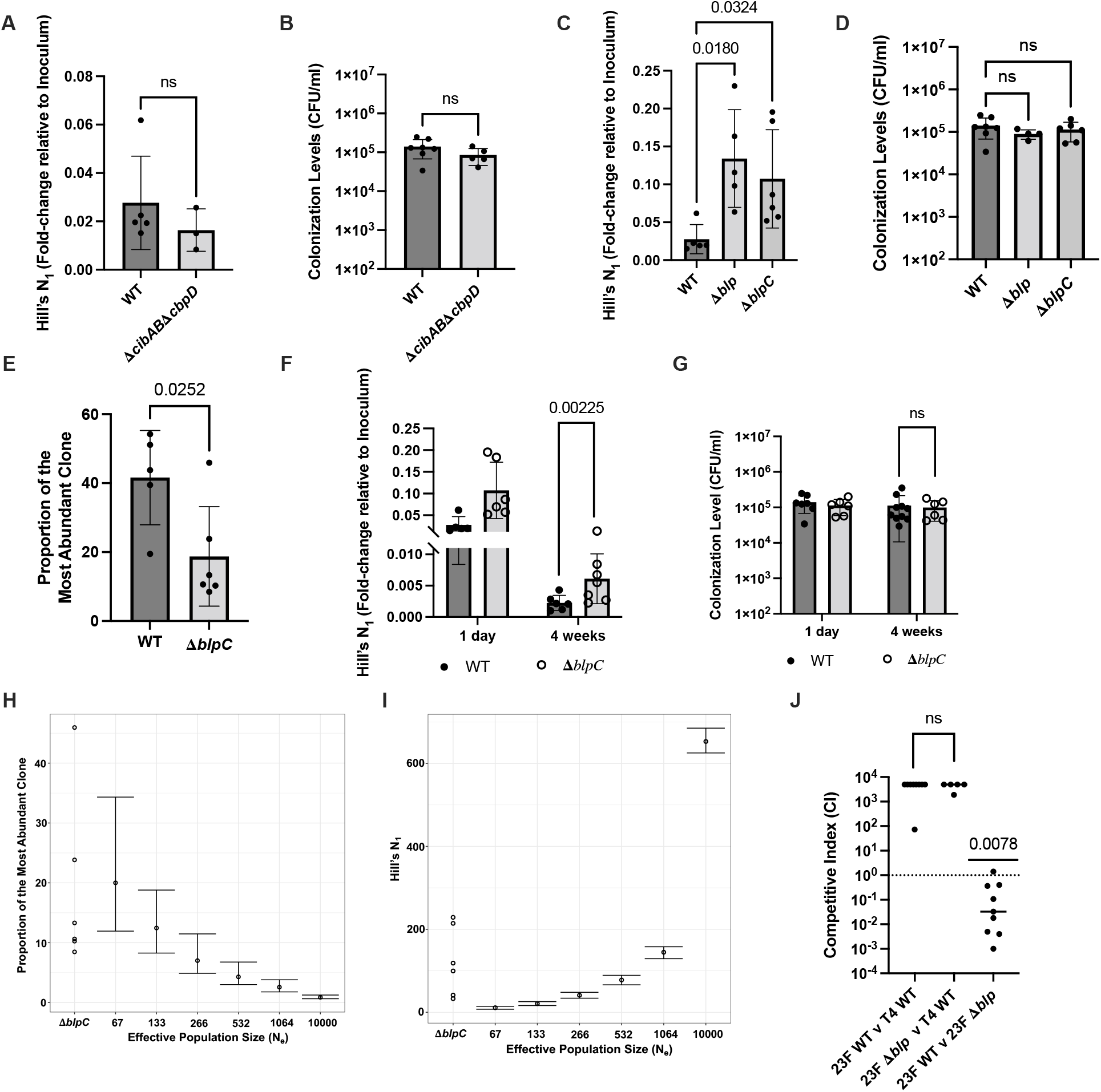
BlpC-mediated signaling results in the *Spn* loss of diversity during colonization. (A, B) Fratricidal molecules CibAB and CbpD do not alter *Spn* population structure. Graphs depicting (A) Hill’s N_1_ diversity coefficient, and (B) colonization levels of WT and Δ*cibAB*Δ*cbpD* strains 1 day post-inoculation. (C-G) BlpC-mediated signaling contributes to loss of clonal diversity. Graphs depicting (C) Hill’s N_1_ diversity coefficient, and (D) colonization levels of WT, Δ*blp* locus, and Δ*blpC* strains 1 day post-inoculation. *P*-values were calculated by ANOVA followed by Tukey’s multiple comparison test. (E) Population structure depicted by comparing proportion of the most abundant clone in WT and Δ*blpC* strains 1 day post-inoculation. (F, G) Changed population structure of Δ*blpC* strain is sustained 4 weeks post-inoculation. Graphs depicting (F) Hill’s N_1_ diversity coefficient, and (G) colonization levels of WT and Δ*blpC* strains at 4 weeks post-inoculation. The 1 day data is re-plotted for illustration. *P*-values were calculated by Student’s *t-*test. (H, I) Contribution of genetic drift in reducing clonal diversity of Δ*blpC* strain cannot be ruled out. Comparison of expected and observed proportion of the most abundant clone 1 day post-inoculation for the Δ*blpC* strain. Δ*blpC* denotes the observed value, while the remaining points denote expected values at different effective population sizes. Points show the mean from 300 repeats, and bars the 95% quantile. (J) Graph denotes median of competitive index (CI) values between competing strains from individual pups 2 weeks post-inoculation. CI was calculated as the ratio of strains recovered from infant pups relative to that in the inoculum. No difference in the CI when either 23F WT or Δ*blp* strain were competing against the T4 WT by Student’s *t*-test. The CI were also compared to a median hypothetical value of 1 (indicating no competition) using Wilcoxon test.

Competitive interactions *in vivo* are thought to be mediated at least in part by the action of bacteriocin peptides that can kill or inhibit the growth of other bacteria. The *Spn* genome encodes numerous putative bacteriocin gene clusters, some of which are activated downstream of quorum sensing pathways (Aggarwal et al., 2020; Javan et al., 2018). Of these, the Blp family of bacteriocins facilitates competition between different *Spn* strains (Dawid et al., 2007; Reichmann and Hakenbeck, 2000; de Saizieu et al., 2000; Valente et al., 2016; Wholey et al., 2016). We investigated whether the bactericidal activity encoded by molecules of the *blp* locus was responsible for loss in

*Spn* clonal diversity using a sufficiently diverse Δ*blp* library (**Fig S6B**). There was approximately a fivefold increase in the clonal diversity of Δ*blp* compared to the WT strain 1 day post-inoculation with no change in the colonization levels (**Figs 4C, D**). These results suggested that the *blp* locus facilitated not just inter-strain but also intra-strain killing, and was responsible for decreasing the *Spn* clonal diversity *in vivo*.

The *blp* locus is induced by signaling via the peptide pheromone encoded by *blpC* (de Saizieu et al., 2000). We, therefore, tested whether loss of BlpC-mediated signaling was sufficient to observe the restoration of clonal diversity as observed in the Δ*blp* strain. There was no change in the colonization density when inoculated with diversely barcoded Δ*blpC* strain (**Figs 4D, S6C**). As with Δ*blp* strain, we observed a similar increase in the clonal diversity when infected with Δ*blpC* relative to the WT strain at 1 day post-inoculation (**Fig 4C**). There was a marked change in the abundance distribution of the clones as well. While the most abundant WT clone was present a frequency of 41.6%, the corresponding frequency for the Δ*blpC* clone was 18.7% 1 day post-inoculation (**Fig 4E**). These results suggested that the activation of BlpC-mediated signaling in a clone conferred a competitive advantage at the expense of its kin following infection. Additionally, they suggested that BlpC activates a “selfish” program that works to ensure clonal lineage survival leading to the genetic element’s own success within the population. The increased *Spn* diversity when colonized by Δ*blpC* strain was sustained over time as evidenced by approximately threefold increase in diversity over WT strain 4 weeks post-inoculation but no change in colonization levels (**Figs 4F, G**).

In regards to the remaining loss of diversity of the Δ*blpC* strain, we cannot rule out the contribution of genetic drift in influencing the population structure (**Fig 4H, I**). To account for the contribution of genetic drift alone, we also estimated the N_e_ by fitting the model to our observed diversity data. The estimated N_e_ for the Δ*blpC* strain (point estimate: 790; 95% Credible Interval (CrI) = 620-900) was much higher than that for the WT strain (point estimate: 33; 95% CrI = 6-76). These data indicate that BlpC-mediated signaling significantly restricts the N_e_, and increases the influence of genetic drift in altering *Spn* population dynamics.

Next, we evaluated the contribution of *blp* bacteriocins to inter- and intra-strain competition. When two *Spn* strains are co-inoculated into the same mouse in a murine colonization model, the 23F strain used in this work outcompeted another commonly used strain, TIGR4 (Abruzzo et al., 2022). The *blp* locus was not responsible for this inter-strain competitive advantage of the 23F strain, as both WT and Δ*blp* strains performed similarly in competition with TIGR4 (**Fig 4J**). However, when WT and Δ*blp* strains both belonging to the 23F genomic background were tested against each other in a competitive infection, the Δ*blp* strain outcompeted the WT strain (**Fig 4J**). This intra-strain competition that resulted in the killing of the *blp*^+^ population suggested there was a fitness cost associated with expression of the *blp* locus. The conservation of a selfish genetic element is inherently intertwined with the host organism’s own success in the population. Accordingly, the expression of *blp* locus ensures its propagation in the population via *Spn* clonal expansion at the expense of isogenic kin (**Figs 1E, 4E**).

Together, these results implicate the selfish nature of the *blp* locus.

### Tight bottlenecks limit *Spn* transmission during both acquisition and establishment of colonization

Next, we questioned the implication of altered population structure during colonization for another important aspect of *Spn* lifecycle – transmission which can be divided into the steps of acquisition by a new host and establishment of colonization. The sequencing of a chromosomally-barcoded *Spn* library provided us with the molecular resolution to answer this question. During transmission, only 11.3% of *Spn* clones present in inoculated index pups transmitted to littermate contact pups, highlighting a population bottleneck during the acquisition stage of transmission (**Fig 5A**). The *Spn* population acquired by the contact pups exhibited significantly reduced clonal diversity relative to index pups (**Fig 5B**). There was no change in colonization levels between index and contact pups that acquired *Spn* (**Fig 5C**). While the frequency of the most abundant clone in the index pups was about 23%, it increased to 75.3% in the contact pups with a different clone succeeding in each pup (**Figs 5D, E**).

**Fig 5:**
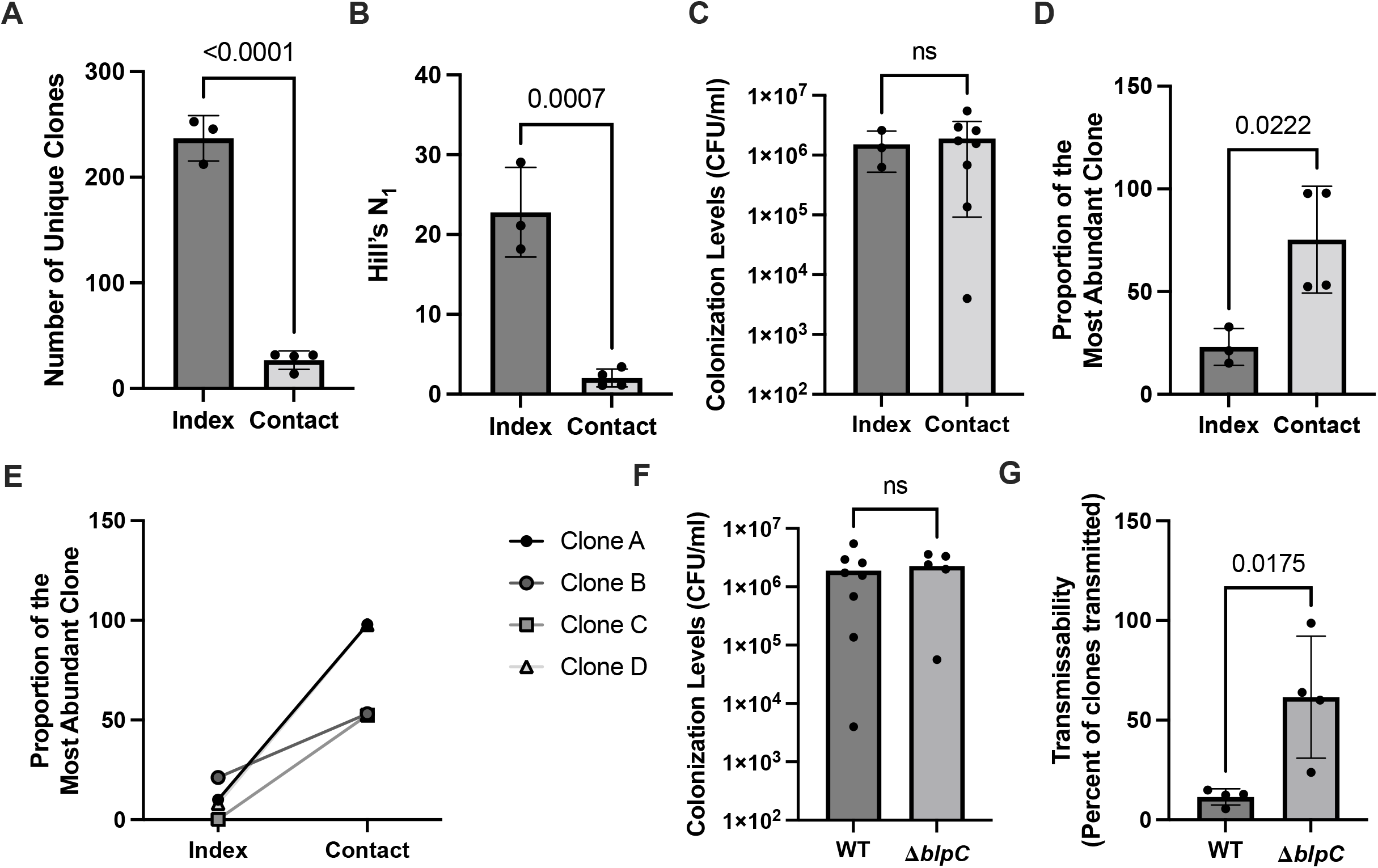
BlpC-mediated signaling reduces the transmissibility of clones. (A-D) Diversity of *Spn* library is significantly reduced in contact pups relative to index pups. Diversity is depicted as (A) number of unique clones, (B) Hill’s N_1_ diversity coefficient, and (D) proportion of the most abundant clone. (C) Graph denotes *Spn* colonization levels in index and contact pups. *P*-values were calculated by Student’s *t*-test. (E) Abundance of clone in the index pup is not necessarily an indicator of its success in establishing colonization in the contact pups. Figure depicts percentage abundance of four clones (A-D) in index and contact pups. Clones were chosen for visualization since these were the most abundant clones in individual contact pups. (F) Colonization levels of WT and Δ*blpC* strains in contact pups. (G) Graph denotes the percentage of clones of WT and Δ*blpC* strains transmitted from index to contact. *P*-values were calculated by Student’s *t*-test.

Following acquisition, not all clones were equally successful in establishing colonization in the contact pups. *Spn* population in the contact pups was dominated by a single clone, pointing towards the existence of another population bottleneck during this stage of transmission. In some cases, clones that were most abundant in the contact pups were present in the index pups at very low frequencies (<0.5%). Our results show that in addition to reduction of clonal diversity during colonization, the clonal pool available for spread within the host population is further restricted during both during acquisition and establishment of *Spn* in a new host. We also attempted to fit bottleneck size to the observed diversity data from our transmission model, obtaining 9 (95% CrI = 1-62), consistent with a bottleneck lower than the fitted N_e_ = 636 for this model, but less than the observed loss of diversity. However, we note that our estimate had a wide credible interval, in part due to uncertainty at which time during the experiment transmission actually occurred.

### Contribution of the *blp* locus to clonal transmission dynamics

Loss of clonal diversity during transmission among infant mice suggested the possibility of a *blp*-mediated effect on transmission success. With this observation in mind, we then evaluated whether data collected from the human population would also support a role of bacteriocins in transmission. We used whole-genome sequence data collected from carriage episodes of unvaccinated mother-child pairs, which has previously been used to identify genetic variation associated with penicillin resistance and carriage duration (Lees et al., 2017). Transmissibility of *Spn* was associated with genetic variations in the *blp* locus (**Table 1**). Although these results were noisy due to the strong correlation between genetic background and phenotype, at least one significant association was found in each *blp* gene. This data hinted at the direct effects of acquired variation within the *blp* locus on transmission rate by conferring fitness advantage for between-host success, or, alternatively could be explained by lineage effects whereby certain *blp* alleles are associated with successfully transmitting clones.

**Table 1:** Top variants in the *blp* genes associated with *Spn* transmissibility.

| **Gene** | **Mutation in D39V\*** | **P-value** |
| --- | --- | --- |
| *blpT* | 476989C->T | 9.33E-08 |
| *blpS* | 477733G->A | 3.57E-14 |
| *blpR* | 477991T->A | 3.10E-17 |
| *blpR* | 478981C->T | 9.62E-24 |
| *blpC* | 479981G->A | 2.96E-16 |
| *blpY* | 484613C->A | 1.65E-20 |
| *blpZ* | 485001T->C | 1.24E-19 |

*\*with reference to the annotated D39V reference genome.*

To directly investigate whether BlpC-mediated signaling played a role in influencing clonal success during transmission, we interrogated how loss of this signaling impacted *Spn* transmission in our murine model. Loss of *blpC* did not influence *Spn* transmissibility (**Fig 5F**). However, there was over a fivefold increase in the percentage of clones acquired by contact pups in the case of Δ*blpC* infection compared to WT *Spn*

(**Fig 5G**). Thus, BlpC-mediated bactericidal activity contributes to population bottlenecks during transmission and colonization. Our data show that an important role of BlpC-mediated bactericidal activity is to limit the pool of clones successfully transmitted to new hosts.

Together, our results highlight the selfish nature of the *blp* locus as it functions to decrease within-host clonal diversity to drive lineage success during transmission to a new host.

## DISCUSSION

An unexplored facet of infection biology is the study of *in vivo* population dynamics that is fundamental for gaining mechanistic insight into bacterial interactions within a host. Here, we investigated this question by focusing on the ecology of colonizing *Spn*. We demonstrate that *Spn* exhibits a rapid loss of clonal diversity early during colonization that results in the eventual domination of the population by a single clonal lineage. This work highlights the non-genetic individuality displayed by isogenic *Spn* population *in vivo*, and how this phenotypic variability results in intra-strain competition by predation of kin cells.

Studies such as omics screens that are aimed at identification of genes important for bacterial colonization and pathogenesis are often noisy. Such screens are typically based on assumptions that the structure of colonizing bacterial populations is homogenous (*e*.*g*., for bulk RNA-seq) and that observed changes in the frequency of a mutant must be due to selection (*e*.*g*., for transposon mutagenesis). However, our results contradict both assumptions by providing evidence of genetic heterogeneity and tight bottlenecks in the colonizing population, as over 70% of the inoculated population was cleared by 1 day post-inoculation. This work highlights the need to account for the effects of intra-strain competition when designing such screens.

Bacterial cells exhibit remarkable spatially resolved transcriptomic heterogeneity during biofilm growth and *in vivo* that can be partly attributed to quorum sensing (Dar et al., 2021; Davis et al., 2015). The BlpC pheromone is an important quorum sensing peptide encoded by *Spn* that signals via BlpHR to exclusively induce genes encoded by the *blp* system (de Saizieu et al., 2000). Activation of BlpC-mediated signaling resulted in the early dominance of a single clone during colonization. One possibility is that following infection, clones that are first to activate the *blp* quorum sensing via inferring local cell density produce bacteriocins and associated immunity proteins. These bacteriocins result in the killing of the neighboring kin cells that have not yet induced the expression of their Blp immunity proteins. Alternatively, stochasticity in bacteriocin-mediated killing during early colonization could increase the representation of a given clone, providing a feedforward advantage.

The *blp* locus, which is under negative frequency-dependent selection, is one of the most heterogenous regions in the *Spn* genome (Bogaardt et al., 2015; Miller et al., 2016). Our results demonstrate that it is not just differences in the possession of cognate immunity proteins but the spatiotemporal dynamics of *blp* activation that leads to decreased *Spn* diversity and increased clonal dominance in a ‘winner takes all’ regime.

These results are striking because they establish that the function of *blp* locus extends beyond inter-strain to intra-strain killing as well. BlpC signaling-mediated killing of isogenic kin results in restricting the N_e_, thereby increasing the influence of genetic drift in shaping the population structure. While bacteriocin-mediated killing of neighboring bacteria increases the amount of free DNA available for uptake by transformation, this may also be detrimental to the bacterial population by increasing the probability of deleterious alleles being fixed in the population. Further, even though the expression of *blp* locus is costly to the cell during competitive interactions *in vivo*, it results in expansion of individual clonal lineages that can ensure the genetic element’s success in the population. This paradox can be explained by considering BlpC-mediated signaling as a selfish program.

Another important aspect of *Spn* lifecycle that is important for its epidemiological success within the host population is the transmission of bacteria between hosts. While there is no stringent bottleneck during the shedding stage (Kono et al., 2016), we show that population bottlenecks exist at the acquisition and colonization stages that limit *Spn* transmission between hosts. Of note, the increased abundance of a clone in the index mouse was not necessarily an indicator of its success during transmission. Our results demonstrate that BlpC-mediated signaling not only limits the pool of clones that are available for transmission, but also restricts their success during spread within the host population. Thus, BlpC-mediated bactericidal activity influences success of a clone both within-and between-hosts.

Analysis of *Spn* isolates recovered from natural human transmission reveal positive associations between genomic variation in the *blp* locus and increased *Spn* transmissibility. We posit that *blp* locus is a determining factor in lineage-dependent success and this is primarily due to intra-rather than inter-strain predation. Genes from the *blp* locus were also found to be frequently mutated during nasopharyngeal carriage in infants (Chaguza et al., 2020). The sustained selective pressure during both colonization and transmission could explain the high genomic heterogeneity of the *blp* locus.

Thus, our findings demonstrate that the rapid loss of clonal diversity observed during *Spn* colonization and transmission is due to a selfish program activated by BlpC. While our current study was focused on *Spn*, we hypothesize that other bacteria may employ similar bet-hedging strategies during infection. It is now increasingly appreciated that various bacterial species routinely employ intra- and inter-strain predation to perpetuate their own survival (Arenas et al., 2015; Bushin et al., 2020; González-Pastor et al., 2003; Jamet et al., 2015; Zhu et al., 2021). Both quorum sensing and bacteriocin production are behaviors commonly exhibited by microbes. It remains to be investigated how the bactericidal molecules encoded by other bacteria also play a role in determining their clonal success *in vivo*.

## MATERIALS & METHODS

### Ethics statement

All animal experiments were performed according to the guidelines laid by National Science Foundation Animal Welfare Act (AWA) and the Public Health Service Policy on the Humane Care and Use of Laboratory Animals. NYU’s Grossman School of Medicine’s Institutional Animal Care and Use Committee (IACUC) oversees the welfare, well-being, proper care, and use of all animals. They have approved the protocols used in this study: IA16-00538 and IA16-00435.

### Bacterial strains and growth conditions

*Spn* 23F strain, P2499, was used in experimental work unless otherwise indicated. P2499 is a streptomycin-resistant derivative of P1121, an ST33 strain obtained from a study of experimental human *Spn* colonization, and previously tested to colonize mice efficiently (Hammond et al., 2021; McCool et al., 2002; Shen et al., 2019; Zafar et al., 2016; Zangari et al., 2021). A mutant lacking the fratricidal genes (Δ*cibAB*Δ*cbpD*, P2576; kanamycin-resistant) was also used in this work (Shen et al., 2019). A streptomycin-resistant derivative of TIGR4, P2406, was used for competition experiments (Zafar et al., 2016). All bacterial strains used in this experimental work are listed in **Table S1**. Colonies were grown from frozen stocks by streaking on TSA-II agar plates supplemented with 5% sheep blood (BD BBL, NJ, USA). Unless otherwise stated, starter cultures were prepared by inoculating streaked colonies in tryptic soy (TS) broth statically at 37°C until they reached an optical density at 620 nm (OD_620_) of 1.0. The cells were then pelleted, washed and resuspended in sterile phosphate-buffered saline (PBS) for mouse inoculations. Bacterial numbers were enumerated by plating serial dilutions on TSA plates supplemented with 100 µl of catalase (38,000 U/ml; Worthington Biochemical Corporation, NJ) and the desired antibiotic (250 µg/ml kanamycin, 200 µg/ml streptomycin, or 200 µg/ml spectinomycin) and incubated overnight at 37°C with 5% CO_2_.

### Construction of mutants

*Spn* mutants were constructed as previously described (Abruzzo et al., 2022; Aggarwal et al., 2021b, 2021a). Colonies were picked and inoculated in acidic Columbia broth (pH 6.6) and grown until an OD_595_ of 0.05 followed by the addition of 5 µg/ml of 1:1 mix of CSP1 and CSP2 along with 500 ng of transforming DNA. Cultures were incubated statically at 37°C with 5% CO_2_ followed by plating on TSA plates supplemented with 100 µl of catalase (38,000 U/ml; Worthington Biochemical Corporation, NJ) and the desired antibiotic (250 µg/ml kanamycin or 200 µg/ml streptomycin). Site-directed homologous recombination was used to create the clean deletion strains, Δ*blp* locus (P2706) and Δ*blpC* (P2646). These mutants were created in a two-step process. In the first step, the *blp* locus (or the *blpC* gene) were replaced with a Janus cassette (containing ∼1kb in flanking regions both upstream and downstream of the region of interest) in P2499 to obtain *blp::*Janus (P2700) and *blpC*::Janus (P2644) constructs. Both P2700 and P2644 were kanamycin resistant. Thereafter, unmarked, clean deletions were made by replacement of the Janus cassette as previously described (Shen et al., 2019; Sung et al., 2001). P2700 and P2644 were transformed with PCR fragments containing ∼1kb each upstream and downstream of the regions of interest joined together to obtain P2706 and P2646, respectively. The desired mutants were streptomycin resistant but kanamycin sensitive. Mutants were confirmed by PCR following each step. All the primers used in this work are listed in **Table S2**.

### Construction of barcoded library

To construct a DNA-barcoded *Spn* library, the designed barcodes were first inserted into a pE539 plasmid vector and cloned into *Escherichia coli*. We designed 7-nt barcodes of the sequence <u>NNM</u>CAATG<u>NNM</u>CAA<u>N</u>, with intervening fixed sequences designed to avoid the presence of start and stop codons. The plasmid pE539 is a derivative of pHSG399 containing a spectinomycin resistance cassette flanked by regions having homology to the *S. pneumoniae* IgA1 protease gene *iga*. The IgA1 protease cleaves human, but not murine, IgA, and so disruption has no known impact on bacterial fitness in mice (Weiser et al., 2003). The barcodes were inserted immediately upstream of the spectinomycin resistance cassette and in the disrupted *iga* gene. The synthesized oligos containing the barcodes were inserted into the plasmid vector by isothermal assembly at 50°C for 1 hour. The assembled plasmid pool was transformed into 5-alpha competent *E. coli* (New England Biolabs, MA) and the transformants were selected by plating on Luria broth (LB) agar plates supplemented with 200 µg/ml spectinomycin. Using this approach, we obtained 3,725 uniquely barcoded *E. coli* cells with the most abundant clone present at a frequency of 0.16% (**Fig S1**). The transformed *E. coli* cells were then grown to isolate the pooled plasmid library by using a Plasmid Midi kit (Qiagen, Germantown, MD) as per manufacturer’s instructions. The transformed *E. coli* library served as a reservoir for uniquely barcoded-plasmids and was stocked at -80°C. The pooled plasmid library was then transformed into *Spn* strains using the protocol described above. The barcoded *Spn* transformants were selected on TS plates supplemented with 100 µl of catalase (38,000 U/ml; Worthington Biochemical Corporation, NJ) and 200 µg/ml spectinomycin. The resulting barcoded *Spn* library was grown, sequenced and stocked at -80°C.

### Library sequencing

Genomic DNA from the samples was isolated using MasterPure Complete DNA & RNA Purification Kit (Lucigen, Middleton, WI) as per manufacturer’s instructions. Barcodes were amplified from genomic DNA using Nested PCR; wherein, the first step consisted of amplifying the *iga* region (5 cycles) followed by amplification of the barcodes (35 cycles). Primers used for amplification of the barcodes contained the adapters to be used for sequencing library preparation. These amplicons were then purified using QIAquick PCR purification kit (Qiagen, Germantown, MD) as per manufacturer’s instructions. Purified samples were then shipped to Azenta Life Sciences (South Plainfield, NJ) for sequencing using their Next-Gen Amplicon-EZ service.

### Analysis of sequencing data

Reads were aligned to a reference sequence using Python. First, trimmomatic was used for quality control to trim adaptor sequences and low-quality bases from the reads (sliding window size: 3, sliding window quality: 20, leading and trailing quality: 15, minimum length: 75) (Bolger et al., 2014). The reads were then aligned to a reference sequence by BWA (Matching Score: 10, Mismatch Penalty: 2) and outputted in a .sam file (Li, 2013). The remainder of the analysis was done using R. The barcode sequence was extracted from the aligned reads by concatenating bases at known variable positions while filtering out incomplete or ambiguous barcodes. A table detailing each barcode detected and the number of times it was found was compiled. To account for variability in the number of total reads, we standardized samples by computing rarefaction and extrapolation of clonal diversity using iNEXT (Chao et al., 2014; Hsieh et al., 2016). The clonal diversity was expressed using Hill numbers with q=0 (‘clonal richness’ or number of unique clones present) and q=1 (Hill’s N_1_) (Hsieh et al., 2016). Shannon diversity index (H) was calculated as *H* = -⅀*p*_*i*_. ln(*p*_*t*_) where *p*_*i*_ denotes the proportion of the population made up of the clone *i*.

### Animal studies

Wild-type C57BL/6J (strain 00664), congenic *Tlr2*^*-/-*^ (strain 004650), and *Ilr1*^*-/-*^ (strain 003245) mice were purchased from The Jackson Laboratory (Bar Harbor, ME). *Il17ra*^*-/-*^ mice were obtained from Amgen (Thousand Oaks, CA). Each of these mouse colonies was bred and maintained in a conventional mouse facility. Infant pups were housed with the dam until weaning at 3 weeks of age. Following infection, all mice appeared healthy, showed normal activity, and gained weight similarly compared to uninfected controls.

Weaned (and adult) mice were fed *ad lib* the PicoLab Rodent Diet 20, a 20% protein diet formulation, and were given water for consumption. All the animals were kept on a light-cycle of 12◻hours on, 12◻hours off with a temperature in the animal facility of 70°F (±2°F).

### Infant and adult colonization model

Three to four day old infant pups were inoculated intranasally with 10^5^ CFU of *Spn* in 3 µl of sterile PBS with a pipette tip, without anesthesia. The pups were returned to their dam for the duration of the experiment. The high dose of *Spn* was used to avoid potentially imposing a population bottleneck during inoculation. For intranasal inoculation of adult mice (6-8 weeks of age), 10^5^ CFU of *Spn* was instilled in 10 ul of sterile PBS. For host microbiota depletion, adult mice were fed with 500 mg/l streptomycin in drinking water three days prior to *Spn* infection. For co-infection with influenza A virus (IAV), mice were inoculated with 250 pfu of IAV/HKx31 in 3 µl of PBS four days after *Spn* infection. At the end of the experiments, mice were euthanized at the indicated time point by CO_2_ asphyxiation followed by cardiac puncture. The *Spn* colonization density of the upper respiratory tract was measured as previously described (Richard et al., 2014). Briefly, the trachea was lavaged using a 30-gauge needle for infants (26-gauge for adults) with 300 µl of sterile PBS collected from the nares. 40 µl of this retrotracheal lavage was used to enumerate bacteria by viable plating serial dilutions on TSA-catalase plates supplemented with the appropriate antibiotic (250 µg/ml kanamycin, 200 µg/ml streptomycin, or 200 µg/ml spectinomycin) and incubated overnight at 37°C with 5% CO_2_. The remaining lavage was grown in TS broth supplemented with 200 µg/ml spectinomycin at 37°C until it reached an OD_620_ of 1.0 for genomic DNA isolation.

### Transmission model

Transmission studies were carried out in infant pups as previously described (Richard et al., 2014). Briefly, one four day-pup per litter (or “index” pup) was intranasally inoculated with 10^5^ CFU of *Spn* in 3 µl of sterile PBS and returned to the litter. The ratio of index to uninfected “contact” pups per litter was 1:3 or 1:4. Four days after *Spn* inoculation, both the index and contact pups were infected with 250 pfu of IAV/HKx31 in 3 µl of PBS. Then, 5-6 days post-IAV infection (9-10 days after *Spn* infection), all pups were euthanized and the bacterial density in the URT of these pups was quantified by viable plating. Transmission was evaluated by testing whether contact pups showed *Spn* colonization.

### Genetic drift estimations

Experimental data observed from pups sacrificed at 6 hours post-inoculation was modelled to investigate whether genetic drift was solely responsible for changes in the population structure observed at 1-day post-inoculation. We performed Wright-Fisher simulations to simulate genetic drift, which allowed us to calculate complex diversity indices, use the empirical starting frequencies in the inoculum to account for founder effects and include stochasticity. The members of each subsequent generation is drawn from a multinomial distribution with probabilities set equal to the frequency in the current generation, such that every clone has equal fitness and a single parent. All generations are of equal size (the effective population size N_e_). At each generation, we calculated the number of unique clones, the maximum abundance of any clone, and the Hill’s N_1_ diversity. Using median colonization levels at 2- and 6 hours post-inoculation, an *in vivo* doubling time of 180 mins was used for calculations. This doubling time amounted to 6 cycles of division between 6 hours and 1 day post-inoculation. For the transmission model, we introduced an extra sample down to the bottleneck size halfway through the simulation, followed by a return to the effective population size.

To fit this model to our experimental data we used sequential approximate Bayesian computation, using Beaumont’s adaptive algorithm (Beaumont et al., 2009). We used the Hill’s N_1_ diversity index and maximum abundance as summary statistics, and uniform priors on the effective population size (between 1 and 2000) and the bottleneck size (between 1 and fitted N_e_).

### CFSE assay

The assay was performed as previously described (Abruzzo et al., 2022). Briefly, *Spn* grown to an OD_620_ of 1.0 were resuspended in 1 ml PBS containing 1% catalase and 10 µM carboxyfluorescein diacetate succinimidyl ester dye (CFSE, Thermo Fisher Scientific, Waltham, MA) and incubated at 37°C for 25 mins. Cells were then washed three times with PBS to remove excess dye. Mice were then inoculated with 10^7^ CFU of CFSE-labeled *Spn* as described above. Immediately following infection (0 hours post-inoculation) and 19 hours post-inoculation, mice were sacrificed and *Spn* in the retrotracheal nasal lavages was analyzed using flow cytometry. The time point of 19 hours was chosen since this was the latest time point where *Spn* replication could be assessed. Samples were fixed by treatment with 2% paraformaldehyde for 25 mins.

They were stained with typing serum specific to 23F (Statens Serum Institut, Copenhagen, Denmark) at 1:1000 dilution by incubating for 25 mins at 4°C. Thereafter, they were incubated with phycoerythrin (PE)-labeled secondary anti-rabbit IgG at 1:100 dilution overnight at 4°C. Flow cytometry was performed using the LSRII flow cytometer (BD Biosciences, Franklin Lakes, NJ) and analyzed by FlowJo Software v10 Software (BD Life Sciences, Ashland, OR). Owing to the even split of CFSE dye between the two daughter cells per replication, a 50% decline in median fluorescence intensity of CFSE in the PE-positive population (*Spn*) was considered a cell division.

### Competition experiment

For competition experiment, strains were grown in TS broth until they reached an OD_620_ of 1.0. The cells were collected, washed and resuspended in sterile PBS. Both the strains were then mixed at a ratio of 1:1. Infant pups were then inoculated intranasally as described above with 10^2^ CFU of the mix. After two weeks, URT lavages were collected and the density of each strain was determine either by selective plating on different antibiotics or by colony immunoblotting using serotype-specific sera (as previously described (Abruzzo et al., 2022)). A competitive index (CI) was calculated by comparing the output CFU ratio of the two strains obtained from the lavages to their input ratio from the inoculum. CI greater than 1 suggests a competitive advantage to the first strain, while a value less than 1 indicates an advantage for the second strain. For pups where only one strain was detected, hypothetical CI values of 1000 were assumed.

### Genomic variations and *Spn* transmissibility

We used whole-genome sequence data collected from carriage episodes of unvaccinated mother-child pairs, which has previously been used to identify genetic variation associated with penicillin resistance and carriage duration. To define a transmissibility phenotype, we divided the dataset into genetic clusters using a k-mer based approach, PopPUNK (Lees et al., 2019). 91 of these clusters had at least four samples. In these clusters, we reconstructed the number of transmission events using transphylo (Didelot et al., 2017), assuming an R0 of 1, sampling rate of 0.12, and average carriage duration of 61 days, as has been previously used for this dataset (Tonkin-Hill et al., 2022). We then divided transmissions per cluster by the time to most recent common ancestor to obtain a rough estimate of transmissibility. We then used pyseer (Lees et al., 2018) to associate SNP variation mapped against the D39V reference (Slager et al., 2018) with this phenotype, while adjusting for population structure.

### Growth and clearance calculations

The changes in allele frequency of the most abundant clone were estimated by simulating *Spn* population during colonization as previously described (Li et al., 2013). We used the same assumption that the colonizing population was comprised of two sub-populations: one small, actively replication sub-population (P_1_) and the other larger sub-population (P_2_) where most bacterial death and clearance occurs. Both sub-populations were assumed to consist of two clones: A (the most abundant clone) and the rest of the clones being grouped together as one (O). The population dynamics of these clones were calculated using the equations as described in that work. The stable equilibrium population was calculated under same assumptions, with a growth rate of 0.0231 min^-1^, clearance rate of 0.0043 min^-1^, and carrying capacities for both the sub-populations as 4000 and 400000. The initial state was calculated by assuming the frequency of an allele in the population to be equivalent to the observed frequency of most abundant clone at 6 hours post-inoculation. Growth and clearance rates were then varied for either of the clones (A or O) to estimate the frequency of the most abundant clone 1 day post-inoculation.

### Statistical analysis

The statistical analyses were performed using GraphPad Prism v9.3.1 (GraphPad Software Inc., San Diego, CA) unless stated otherwise.

## Code availability

The code for analyzing the sequencing data is available at https://github.com/sda26/pneumo_diversity. The code for running population genetic simulations, fitting to data, and fitting generalized linear models is available at https://github.com/johnlees/transmission_blp (version 1.0, Apache 2.0 license).

## Acknowledgements

We thank Dr. Kristen Lokken-Toyli for her suggestions and feedback. The illustrations were created with BioRender.com. This work was funded by NIH grants R01 AI50893, R01 AI038446, and R21 AI50867 to JNW. NTJ was supported by NIH grant T32 AI007180-37.

## REFERENCES

Abruzzo, A.R., Aggarwal, S.D., Sharp, M.E., Bee, G.C.W., and Weiser, J.N. (2022). Serotype-dependent effects on the dynamics of pneumococcal colonization and implications for transmission. MBio 13, e00158–22. https://doi.org/10.1128/mbio.00158-22.

Aggarwal, S.D., Yesilkaya, H., Dawid, S., and Hiller, N.L. (2020). The pneumococcal social network. PLoS Pathog. 16, e1008931. https://doi.org/https://doi.org/10.1371/journal.ppat.1008931.

Aggarwal, S.D., Gullett, J.M., Fedder, T., Safi, J.P.F., Rock, C.O., and Hiller, N.L. (2021a). Competence-associated peptide BriC alters fatty acid biosynthesis in Streptococcus pneumoniae. MSphere 6, e00145–21.

Aggarwal, S.D., Lloyd, A.J., Yerneni, S.S., Narciso, A.R., Shepherd, J., Roper, D.I., Dowson, C.G., Filipe, S.R., and Hiller, N.L. (2021b). A molecular link between cell wall biosynthesis, translation fidelity, and stringent response in Streptococcus pneumoniae. Proc. Natl. Acad. Sci. 118, e2018089118. https://doi.org/10.1073/pnas.2018089118/-/DCSupplemental.Published.

Arenas, J., De Maat, V., Catón, L., Krekorian, M., Herrero, J.C., Ferrara, F., and Tommassen, J. (2015). Fratricide activity of MafB protein of N. meningitidis strain B16B6. BMC Microbiol. 15, 156. https://doi.org/10.1186/s12866-015-0493-6.

Bachta, K.E.R., Allen, J.P., Cheung, B.H., Chiu, C.H., and Hauser, A.R. (2020). Systemic infection facilitates transmission of Pseudomonas aeruginosa in mice. Nat. Commun. 11, 543. https://doi.org/10.1038/s41467-020-14363-4.

Beaumont, M.A., Cornuet, J.-M., Marin, J.-M., and Robert, C.P. (2009). Adaptive approximate Bayesian computation. Biometrika 96, 983–990. https://doi.org/10.1093/biomet/asp052.

Bogaardt, C., van Tonder, A.J., and Brueggemann, A.B. (2015). Genomic analyses of pneumococci reveal a wide diversity of bacteriocins - including pneumocyclicin, a novel circular bacteriocin. BMC Genomics 16, 554. https://doi.org/10.1186/s12864-015-1729-4.

Bolger, A.M., Lohse, M., and Usadel, B. (2014). Trimmomatic: A flexible trimmer for Illumina sequence data. Bioinformatics 30, 2114–2120. https://doi.org/10.1093/bioinformatics/btu170.

Bushin, L.B., Covington, B.C., Rued, B.E., Federle, M.J., and Seyedsayamdost, M.R. (2020). Discovery and biosynthesis of Streptosactin, a sactipeptide with an alternative topology encoded by commensal bacteria in the human microbiome. J. Am. Chem. Soc. 142, 16265–16275. https://doi.org/10.1021/jacs.0c05546.

Chaguza, C., Senghore, M., Bojang, E., Gladstone, R.A., Lo, S.W., Tientcheu, P.E., Bancroft, R.E., Worwui, A., Foster-Nyarko, E., Ceesay, F., et al. (2020). Within-host microevolution of Streptococcus pneumoniae is rapid and adaptive during natural colonisation. Nat. Commun. 11, 3442. https://doi.org/10.1038/s41467-020-17327-w.

Chao, A., Gotelli, N.J., Hsieh, T.C., Sander, E.L., Ma, K.H., Colwell, R.K., and Ellison, A.M. (2014). Rarefaction and extrapolation with Hill numbers: A framework for sampling and estimation in species diversity studies. Ecol. Monogr. 84, 45–67. https://doi.org/10.1890/13-0133.1.

D’Mello, A., Riegler, A.N., Martínez, E., Beno, S.M., Ricketts, T.D., Foxman, E.F., Orihuela, C.J., and Tettelin, H. (2020). An in vivo atlas of host–pathogen transcriptomes during Streptococcus pneumoniae colonization and disease. Proc. Natl. Acad. Sci. U. S. A. 117, 33507–33518. https://doi.org/10.1073/pnas.2010428117.

Dar, D., Dar, N., Cai, L., and Newman, D.K. (2021). Spatial transcriptomics of planktonic and sessile bacterial populations at single-cell resolution. Science 373, eabi4882. https://doi.org/10.1126/science.abi4882.

Davis, K.M., Mohammadi, S., and Isberg, R.R. (2015). Community behavior and spatial regulation within a bacterial microcolony in deep tissue sites serves to protect against host attack. Cell Host Microbe 17, 21–31. https://doi.org/10.1016/j.chom.2014.11.008.

Dawid, S., Roche, A.M., and Weiser, J.N. (2007). The blp bacteriocins of Streptococcus pneumoniae mediate intraspecies competition both in vitro and in vivo. Infect. Immun. 75, 443–451. https://doi.org/10.1128/IAI.01775-05.

Didelot, X., Fraser, C., Gardy, J., and Colijn, C. (2017). Genomic infectious disease epidemiology in partially sampled and ongoing outbreaks. Mol. Biol. Evol. 34, 997–1007. https://doi.org/10.1093/molbev/msw275.

Fiebig, A., Vrentas, C.E., Le, T., Huebner, M., Boggiatto, P.M., Olsen, S.C., and Crosson, S. (2021). Quantification of Brucella abortus population structure in a natural host. Proc. Natl. Acad. Sci. U. S. A. 118, e2023500118. https://doi.org/10.1073/pnas.2023500118.

González-Pastor, J.E., Hobbs, E.C., and Losick, R. (2003). Cannibalism by sporulating bacteria. Science 301, 510–513. https://doi.org/10.1126/science.1086462.

Hammond, A.J., Binsker, U., Aggarwal, S.D., Ortigoza, M.B., Loomis, C., and Weiser, J.N. (2021). Neuraminidase B controls neuraminidase A-dependent mucus production and evasion. PLoS Pathog. 17, e1009158. https://doi.org/https://doi.org/10.1371/ journal.ppat.1009158 Editor:

Hsieh, T.C., Ma, K.H., and Chao, A. (2016). iNEXT: an R package for rarefaction and extrapolation of species diversity (Hill numbers). Methods Ecol. Evol. 7, 1451–1456. https://doi.org/10.1111/2041-210X.12613.

Hullahalli, K., and Waldor, M.K. (2021). Pathogen clonal expansion underlies multiorgan dissemination and organ-specific outcomes during systemic infection. Elife 10, e70910..

Hullahalli, K., Pritchard, J.R., and Waldor, M.K. (2021). Refined quantification of infection bottlenecks and pathogen dissemination with STAMPR. MSystems 6, e00887–21.

Jamet, A., Jousset, A.B., Euphrasie, D., Mukorako, P., Boucharlat, A., Ducousso, A., Charbit, A., and Nassif, X. (2015). A new family of secreted toxins in pathogenic Neisseria species. PLoS Pathog. 11, e1004592. https://doi.org/10.1371/journal.ppat.1004592.

Jasinska, W., Manhart, M., Lerner, J., Gauthier, L., Serohijos, A.W.R., and Bershtein, S. (2020). Chromosomal barcoding of E. coli populations reveals lineage diversity dynamics at high resolution. Nat. Ecol. Evol. 4, 437–452.

Javan, R.R., van Tonder, A.J., King, J.P., Harrold, C.L., and Brueggemann, A.B. (2018). Genome sequencing reveals a large and diverse repertoire of antimicrobial peptides. Front. Microbiol. 9, 2012. https://doi.org/10.3389/fmicb.2018.02012.

Kono, M., Zafar, M.A., Zuniga, M., Roche, A.M., Hamaguchi, S., and Weiser, J.N. (2016). Single cell bottlenecks in the pathogenesis of Streptococcus pneumoniae. PLoS Pathog. 12, e1005887. https://doi.org/10.1371/journal.ppat.1005887.

Kuipers, K., Lokken, K.L., Zangari, T., Boyer, M.A., Shin, S., and Weiser, J.N. (2018). Age-related differences in IL-1 signaling and capsule serotype affect persistence of Streptococcus pneumoniae colonization. PLoS Pathog. 14, e1007396. https://doi.org/10.1371/journal.ppat.1007396.

Lees, J.A., Croucher, N.J., Goldblatt, D., Nosten, F., Parkhill, J., Turner, C., Turner, P., and Bentley, S.D. (2017). Genome-wide identification of lineage and locus specific variation associated with pneumococcal carriage duration. Elife 6, e26255. https://doi.org/10.7554/eLife.26255.

Lees, J.A., Galardini, M., Bentley, S.D., Weiser, J.N., and Corander, J. (2018). pyseer: A comprehensive tool for microbial pangenome-wide association studies. Bioinformatics 34, 4310–4312. https://doi.org/10.1093/bioinformatics/bty539.

Lees, J.A., Harris, S.R., Tonkin-Hill, G., Gladstone, R.A., Lo, S.W., Weiser, J.N., Corander, J., Bentley, S.D., and Croucher, N.J. (2019). Fast and flexible bacterial genomic epidemiology with PopPUNK. Genome Res. 29, 304–316. https://doi.org/10.1101/gr.241455.118.

Li, H. (2013). Aligning sequence reads, clone sequences and assembly contigs with BWA-MEM. 1303.3997v2.

Li, Y., Thompson, C.M., Trzciński, K., and Lipsitch, M. (2013). Within-host selection is limited by an effective population of Streptococcus pneumoniae during nasopharyngeal colonization. Infect. Immun. 81, 4534–4543. https://doi.org/10.1128/IAI.00527-13.

Lu, Y.-J., Gross, J., Bogaert, D., Finn, A., Bagrade, L., Zhang, Q., Kolls, J.K., Srivastava, A., Lundgren, A., Forte, S., et al. (2008). Interleukin-17A mediates acquired immunity to pneumococcal colonization. PLoS Pathog. 4, e1000159. https://doi.org/10.1371/journal.ppat.1000159.

Martin, C.J., Cadena, A.M., Leung, V.W., Lin, P.L., Maiello, P., Hicks, N., Chase, M.R., Flynn, J.A.L., and Fortune, S.M. (2017). Digitally barcoding Mycobacterium tuberculosis reveals In vivo infection dynamics in the macaque model of tuberculosis. MBio 8, e00312–17. https://doi.org/10.1128/mBio.00312-17.

Matthews, A.J., Rowe, H.M., Rosch, J.W., and Camilli, A. (2021). A Tn-seq screen of Streptococcus pneumoniae uncovers DNA repair as the major pathway for desiccation tolerance and transmission. Infect. Immun. 89, e00713–20. https://doi.org/10.1128/IAI.00713-20.

McCool, T.L., Cate, T.R., Moy, G., and Weiser, J.N. (2002). The immune response to pneumococcal proteins during experimental human carriage. J. Exp. Med. 195, 359–365. https://doi.org/10.1084/jem.20011576.

Miller, E.L., Abrudan, M.I., Roberts, I.S., and Rozen, D.E. (2016). Diverse ecological strategies are encoded by Streptococcus pneumoniae bacteriocin-like peptides. Genome Biol. Evol. 8, 1072–1090. https://doi.org/10.1093/gbe/evw055.

van Opijnen, T., and Camilli, A. (2012). A fine scale phenotype-genotype virulence map of a bacterial pathogen. Genome Res. 22, 2541–2551. https://doi.org/10.1101/gr.137430.112.22.

Reichmann, P., and Hakenbeck, R. (2000). Allelic variation in a peptide-inducible two-component system of Streptococcus pneumoniae. FEMS Microbiol. Lett. 190, 231–236. https://doi.org/10.1016/S0378-1097(00)00340-2.

Richard, A.L., Siegel, S.J., Erikson, J., and Weiser, J.N. (2014). TLR2 signaling decreases transmission of Streptococcus pneumoniae by limiting bacterial shedding in an infant mouse Influenza A co-infection model. PLoS Pathog. 10, e1004339. https://doi.org/10.1371/journal.ppat.1004339.

van Rossum, A.M.C., Lysenko, E.S., and Weiser, J.N. (2005). Host and bacterial factors contributing to the clearance of colonization by Streptococcus pneumoniae in a murine model. Infect. Immun. 73, 7718–7726. https://doi.org/10.1128/IAI.73.11.7718-7726.2005.

Rowe, H.M., Karlsson, E., Echlin, H., Chang, T.-C.C., Wang, L., van Opijnen, T., Pounds, S.B., Schultz-Cherry, S., and Rosch, J.W. (2019). Bacterial factors required for transmission of Streptococcus pneumoniae in mammalian hosts. Cell Host Microbe 25, 884–891. https://doi.org/10.1016/j.chom.2019.04.012.

de Saizieu, A., Gardes, C., Flint, N., Wagner, C., Kamber, M., Mitchell, T.J., Keck, W., Amrein, K.E., and Lange, R. (2000). Microarray-based identification of a novel Streptococcus pneumoniae regulon controlled by an autoinduced peptide. J. Bacteriol. 182, 4696–4703. https://doi.org/10.1128/JB.182.17.4696-4703.2000.

Shen, P., Lees, J.A., Bee, G.C.W., Brown, S.P., and Weiser, J.N. (2019). Pneumococcal quorum sensing drives an asymmetric owner-intruder competitive strategy during carriage via the competence regulon. Nat. Microbiol. 4, 198–208. https://doi.org/10.1016/j.physbeh.2017.03.040.

Siegel, S.J., Roche, A.M., and Weiser, J.N. (2014). Influenza promotes pneumococcal growth during coinfection by providing host sialylated substrates as a nutrient source. Cell Host Microbe 16, 55–67. https://doi.org/10.1016/j.chom.2014.06.005.

Siegel, S.J., Tamashiro, E., and Weiser, J.N. (2015). Clearance of pneumococcal colonization in infants Is delayed through altered macrophage trafficking. PLoS Pathog. 11, e1005004. https://doi.org/10.1371/journal.ppat.1005004.

Slager, J., Aprianto, R., and Veening, J.-W. (2018). Deep genome annotation of the opportunistic human pathogen Streptococcus pneumoniae D39. Nucleic Acids Res. 46, 9971–9989. https://doi.org/10.1101/283663.

Sung, C.K., Li, H., Claverys, J.P., and Morrison, D.A. (2001). An rpsL cassette, Janus, for gene replacement through negative selection in Streptococcus pneumoniae. Appl. Environ. Microbiol. 67, 5190–5196. https://doi.org/10.1128/aem.67.11.5190-5196.2001.

Tonkin-Hill, G., Ling, C., Chaguza, C., Salter, S.J., Hinfonthong, P., Nikolaou, E., Tate, N., Pastusiak, A., Turner, C., Chewapreecha, C., et al. (2022). Pneumococcal within-host diversity during colonisation, transmission and treatment. BioRxiv 2022.02.20.480002.

Valente, C., Dawid, S., Pinto, F.R., Hinds, J., Simões, A.S., Gould, K.A., Mendes, L.A., de Lencastre, H., and Sá-Leão, R. (2016). The blp locus of Streptococcus pneumoniae plays a limited role in the selection of strains that can cocolonize the human nasopharynx. Appl. Environ. Microbiol. 82, 5206–5215. https://doi.org/10.1128/AEM.01048-16.

Vasquez, K.S., Willis, L., Cira, N.J., Ng, K.M., Pedro, M.F., Aranda-Díaz, A., Rajendram, M., Yu, F.B., Higginbottom, S.K., Neff, N., et al. (2021). Quantifying rapid bacterial evolution and transmission within the mouse intestine. Cell Host Microbe 29, 1454–1468. https://doi.org/10.1016/j.chom.2021.08.003.

Weiser, J.N., Bae, D., Fasching, C., Scamurra, R.W., Ratner, A.J., and Janoff, E.N. (2003). Antibody-enhanced pneumococcal adherence requires IgA1 protease. Proc. Natl. Acad. Sci. U. S. A. 100, 4215–4220. https://doi.org/10.1073/pnas.0637469100.

Weiser, J.N., Ferreira, D.M., and Paton, J.C. (2018). Streptococcus pneumoniae: Transmission, colonization and invasion. Nat. Rev. Microbiol. 16, 355–367. https://doi.org/10.1038/s41579-018-0001-8.

Wholey, W.-Y., Kochan, T.J., Storck, D.N., and Dawid, S. (2016). Coordinated bacteriocin expression and competence in Streptococcus pneumoniae contributes to genetic adaptation through neighbor predation. PLOS Pathog. 12, e1005413. https://doi.org/10.1371/journal.ppat.1005413.

Zafar, M.A., Kono, M., Wang, Y., Zangari, T., and Weiser, J.N. (2016). Infant mouse model for the study of shedding and transmission during Streptococcus pneumoniae monoinfection. Infect. Immun. 84, 2714–2722. https://doi.org/10.1128/IAI.00416-16.

Zafar, M.A., Hammond, A.J., Hamaguchi, S., Wu, W., Kono, M., Zhao, L., and Weiser, J.N. (2019). Identification of pneumococcal factors affecting pneumococcal shedding shows that the dlt locus promotes inflammation and transmission. MBio 10, e01032–19. https://doi.org/10.1128/mBio.01032-19.

Zangari, T., Ortigoza, M.B., Lokken-Toyli, K.L., and Weiser, J.N. (2021). Type I interferon signaling is a common factor driving Streptococcus pneumoniae and Influenza A virus shedding and transmission. MBio 12, e03589–20. https://doi.org/10.1128/mBio.03589-20.

Zhang, Z., Clarke, T.B., and Weiser, J.N. (2009). Cellular effectors mediating Th17-dependent clearance of pneumococcal colonization in mice. J. Clin. Invest. 119, 1899–1909. https://doi.org/10.1172/JCI36731.The.

Zhu, Y., Ma, J., Zhang, Y., Zhong, X., Bai, Q., Dong, W., Pan, Z., Liu, G., Zhang, C., and Yao, H. (2021). CrfP, a fratricide protein, contributes to natural transformation in Streptococcus suis. Vet. Res. 52, 50. https://doi.org/10.1186/s13567-021-00917-x.

