## Supplemental Figures for "BlpC-mediated selfish program leads to rapid loss of *Streptococcus pneumoniae* clonal diversity during infection"

### Fig S1

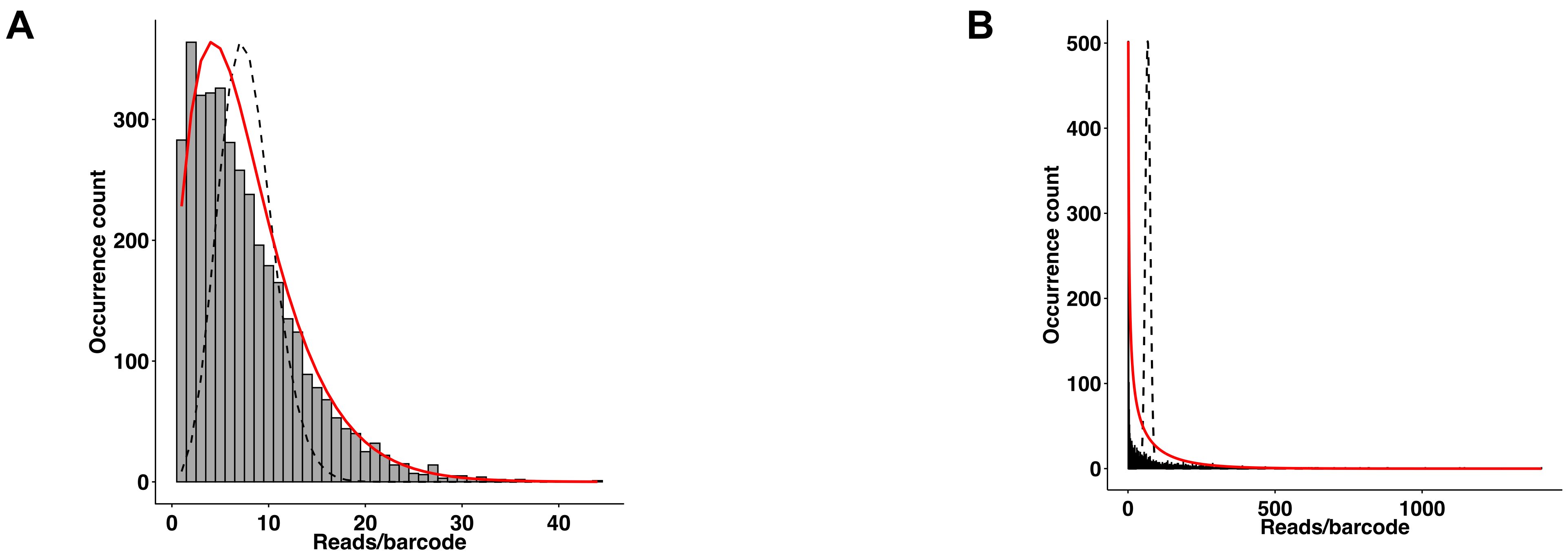


**Fig S1: Molecularly-barcoded clonal library is diverse.** Histograms denote the frequency occurrence (Y-axis) of the number of reads per barcode (X-axis) detected for **(A)** pE539 plasmid, and **(B)** *Spn* libraries. **(A)** The pE539 library contains 3,725 unique barcodes with the proportion of the most abundant barcode present being 0.16%. **(B)** The *Spn* library contains 2,764 uniquely barcoded clones with the proportion of most abundant clone being 0.65%. Both libraries follow a slightly overdispersed Poisson distribution with the following mean and variance values, respectively: **(A)** 7.72 and 31.2, **(B)** 67.68 and 9813.6 . Black dashed line approximates Poisson distribution while the red solid line approximates negative binomial distribution.

### Fig S2

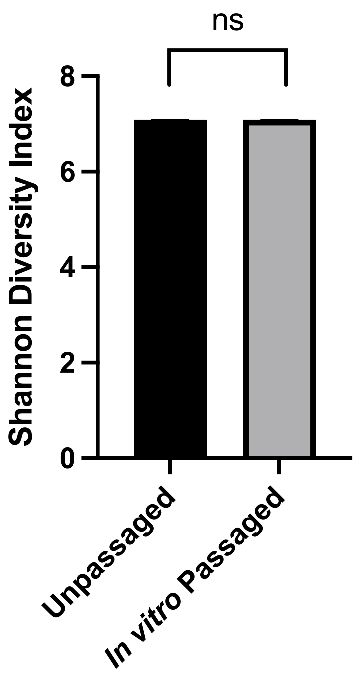


**Fig S2: *Spn* library is stable upon passaging *in vitro***. The unpassaged *Spn* library was inoculated and re-grown in TS broth. Diversity of the *in vitro* passaged library as measured by Shannon Diversity Index (Y-axis) is not different than that of the unpassaged *Spn* library. Error bars denote 95% confidence intervals. *(P>0.05* by Hutcheson's *t*-test*)*

### Fig S3

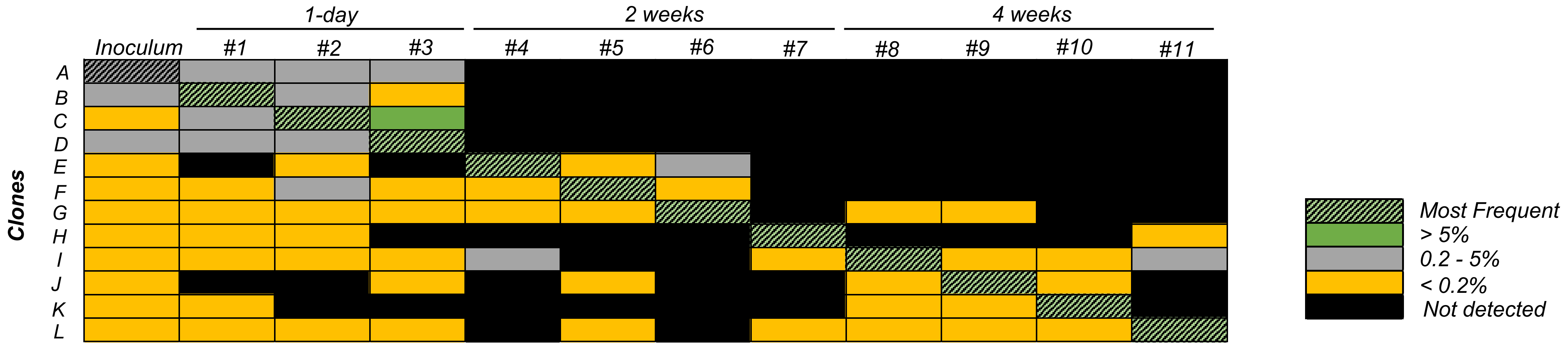


**Fig S3: Success of clones during colonization not due to any fitness difference among cells in the inoculum.** Heat map denotes the abundance of certain selected individual clones that make up the library population either in inoculum or individual mice at different time points. In each of the individual mice, a different clone is detected as being the most abundant clone (hatched) in the population.

**Fig S4**

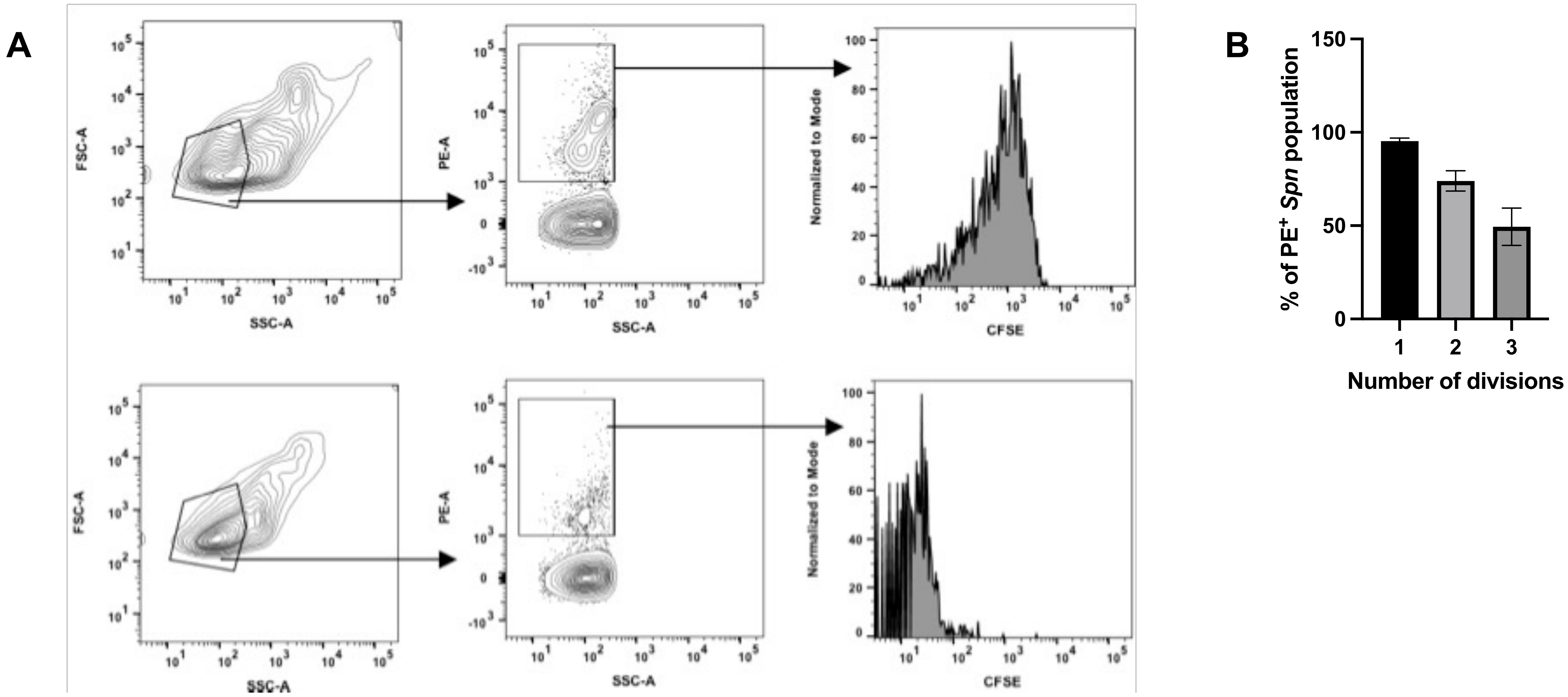


**Fig S4: Majority of *Spn* population is actively replicating during colonization.** Flow cytometry was performed on *Spn* lavaged from the upper respiratory tract of infant mice at 19 hours post-inoculation. *Spn* was identified by staining with 23F-typing sera followed by a phycoerythrin (PE)-secondary antibody. **(A)** Figure denotes gating strategy for CFSE analysis of *Spn* proliferation *in vivo*. *Spn* inoculum pre-stained with CFSE (Top) and with DMSO vehicle control (bottom). **(B)** Graph denotes the percentage of *Spn* population that has undergone 1, 2, or 3 divisions.

### Fig S5

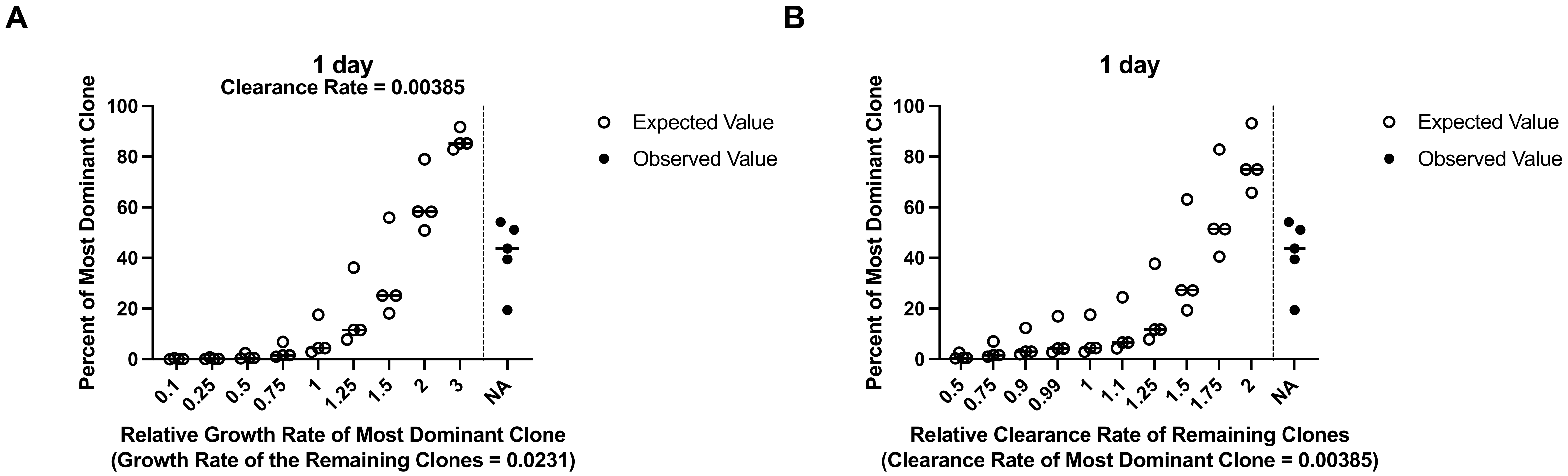


**Fig S5: Variations in growth and clearance rate of individual clones may result in altered population structure.** Expected changed in population structure measured as the expected abundance of the most dominant clone by varying the (A) growth rate of the most dominant clone relative to remaining clones in the population; and (B) clearance rate of the remaining clones relative to the most dominant clone in the population. “NA” corresponds to the experimentally observed values at 1-day post inoculation.

### Fig S6

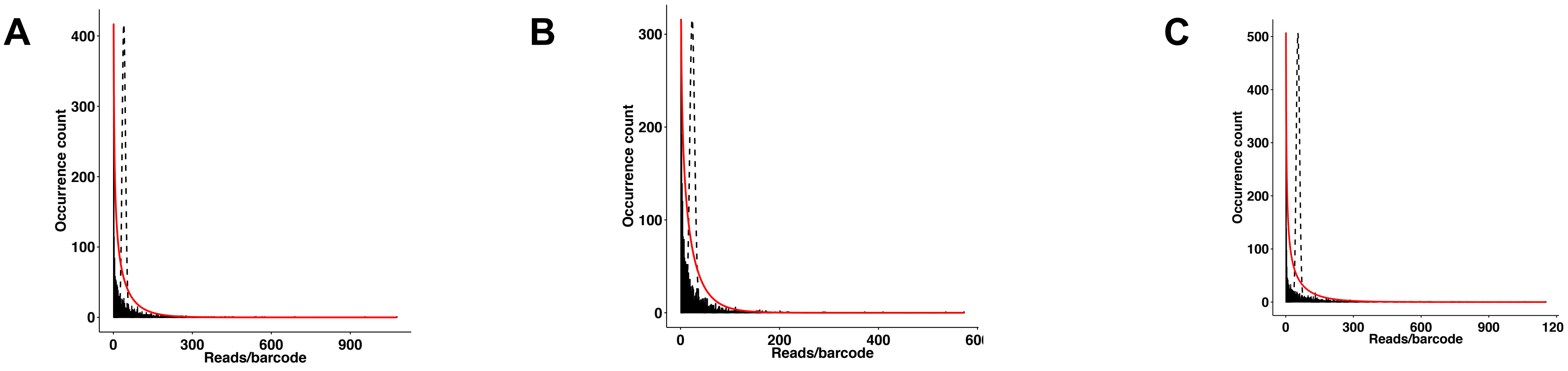


**Fig S6: Molecularly-barcoded libraries in *Spn* mutants are diverse**. Molecularly-barcoded libraries in *Spn* (A) **Δ***cibAB***Δ***cbpD*, (B) **Δ***blp* locus, and (C) **Δ***blpC* strains are diverse. (A) **Δ***cibAB***Δ***cbpD* library contains 2,775 uniquely barcoded clones with the most dominant clone at an abundance of 0.97%. (B) **Δ***blp* locus library contains 2,339 uniquely barcoded clones with the most dominant clone at an abundance of 0.99%. (C) **Δ***blpC* library contains 3,007 uniquely barcoded clones with the most dominant clone at an abundance of 0.7%. All libraries follow slightly overdispersed Poisson distribution with the following mean and variance values, respectively: **(A)** 40.1 and 2861.17, **(B)** 23.99 and 848.82, and **(C)** 54.44 and 6228.84. Black dashed line denotes Poisson distribution and the red solid line denotes negative binomial distribution.
