## Supplemental Table 1 for "BlpC-mediated selfish program leads to rapid loss of *Streptococcus pneumoniae* clonal diversity during infection"

**Table S1: Bacterial strains used in this experimental work**

| Strain | Strain Number | Description | Reference |
|---|---|---|---|
| 23F | P2499 | Streptomycin resistant derivative of 23F (23F WT) | Shen et al (2019) |
| T4 | P2406 | Streptomycin resistant derivative of TIGR4 (T4 WT) | Zafar *et al* (2016) |
| Δ*cbpD* Δ*cibABC* | P2576 | P2499 with clean deletion of *cbpD* and Janus cassette replacing *cibABC* operon, kanamycin resistant | Shen *et al* (2019) |
| *blpC::Janus* | P2644 | P2499 with Janus cassette replacing *blpC* , kanamycin resistant | This study |
| Δ*blpC* | P2646 | P2644 with clean deletion of *blpC,* streptomycin resistant | This study |
| *blp::Janus* | P2700 | P2499 with Janus cassette replacing *blp* locus, kanamycin resistant | This study |
| Δ*blp* | P2706 | P2700 with clean deletion of *blp,* streptomycin resistant | This study |
