## Supplemental Table 2 for "BlpC-mediated selfish program leads to rapid loss of *Streptococcus pneumoniae* clonal diversity during infection"

**Table S2: Primers used in this study**

| Purpose | Primer Name | Primer Number | Sequence |
|---|---|---|---|
| Construction of *blpC::Janus* strain | blpC_Flank1_F | SDA01 | ATACCTTTATGAATTTTACTAAACAATCCCATG |
| | blpC_Flank1_R | SDA02 | CACATTATCCATTAAAAATCAAACGGACTTATCCATG TTATGATTCTCCTTTTTGAT |
| | blpC_Janus_F | SDA03 | ATCAAAAAGGAGAATCATAACATGGATAAGTCCGTTT GATTTTTAATGGATAATGTG |
| | blpC_Janus_R | SDA04 | TTCTCGGTCTTGTTTTTATAGCTGCCTTTCCTTATGCT TTTGGACGTTTAG |
| | blpC_Flank2_F | SDA05 | CTAAACGTCCAAAAGCATAAGGAAAGGCAGCTATAA AAACAAGACCGAGAA |
| | blpC_Flank2_R | SDA09 | TCTCTTACTAAGATTAACTGGGCAATG |
| Construction of Δ*blpC* strain | blpC_Flank1_F | SDA01 | ATACCTTTATGAATTTTACTAAACAATCCCATG |
| | blpC_Clean_R | SDA07 | TCGGTCTTGTTTTTATAGCTGAATAGGTTGTTTCTTAT CCATGTTATGATTCTCCTT |
| | blpC_Clean_F | SDA08 | AAGGAGAATCATAACATGGATAAGAAACAACCTATT CAGCTATAAAAACAAGACCGA |
| | blpC_Flank2_R | SDA09 | TCTCTTACTAAGATTAACTGGGCAATG |
| Construction of *blp::Janus* strain | blp_Flank1_F | SDA52 | ATGAAAGACTTGTTTTTAAAGAGAAAG |
| | blp_Flank1_R | SDA53 | TAAAAATCAAACGGATTAGTCCTGCATCTGACG |
| | blp_Janus_F | SDA54 | CAGATGCAGGACTAATCCGTTTGATTTTTAATGGATA ATG |
| | blp_Janus_R | SDA55 | ACCTTTTCATAATTTCCTTTCCTTATGCTTTTGG |
| | blp_Flank2_F | SDA56 | AAGCATAAGGAAAGGAAATTATGAAAAGGTAAGCAG AAG |
| | blp_Flank2_R | SDA57 | GATTTGCCTCTAGTAATTCTG |
| Construction of Δ*blp* strain | blp_Flank1_F | SDA52 | ATGAAAGACTTGTTTTTAAAGAGAAAG |
| | blp_Clean_R | SDA58 | ACCTTTTCATAATTTTTAGTCCTGCATCTGACG |
| | blp_Clean_F | SDA59 | CAGATGCAGGACTAAAAATTATGAAAAGGTAAGCAG AAG |
| | blp_Flank2_R | SDA57 | GATTTGCCTCTAGTAATTCTG |
| Linearization of pE539 | pE539_F | N/A | AACTAATAACGTAACGTGACTGGC |
| | pE539_R | N/A | GTTGTTTTGTAGGAGTGGCTGCTG |
| Barcoding oligomers | Barcoding_F | N/A | AAAAATCAGCAGCCACTCCTACAAAACAACTGNNM CAATGNNMCAANAACTAATAACGTAACGTGACTGG CAAGAGA |
| | Barcoding_R | N/A | TCTCTTGCCAGTCACGTTACGTTATTAGTTNTTGKNN CATTGKNNCAGTTGTTTTGTAGGAGTGGCTGCTGATT TTT |
| Amplicon Sequencing | IgG_Nested_F | SDA18 | GAAAGCGCATCTCAATTTTAAGACT |
| | IgG_Nested_R | SDA19 | CCCTATGTTCTACACCAGTCTCA |
| | Seq_Adapt_F | SDA30 | ACACTCTTTCCCTACACGACGCTCTTCCGATCTGAG ATTTCAGGTTACACTTATATTGGA |
| | Seq_Adapt_R | SDA31 | GACTGGAGTTCAGACGTGTGCTCTTCCGATCTTTCG TATGTATTCAAATATATCCTCCTC |
